# Analysis of whole genome-transcriptomic organization in brain to identify genes associated with alcoholism

**DOI:** 10.1101/500439

**Authors:** Manav Kapoor, Jen-Chyong Wang, Sean P. Farris, Yunlong Liu, Jeanette McClintick, Ishaan Gupta, Jacquelyn L. Meyers, Sarah Bertelsen, Michael Chao, John Nurnberger, Jay Tischfield, Oscar Harari, Li Zeran, Victor Hesselbrock, Lance Bauer, Towfique Raj, Bernice Porjesz, Arpana Agrawal, Tatiana Foroud, Howard J. Edenberg, R. Dayne Mayfield, Alison Goate

## Abstract

Alcohol exposure triggers changes in gene expression and biological pathways in human brain. We explored alterations in gene expression in the Pre-Frontal Cortex (PFC) of 65 alcoholics and 73 controls of European descent, and identified 129 genes that showed altered expression (FDR < 0.05) in subjects with alcohol dependence. Differentially expressed genes were enriched for pathways related to interferon signaling and Growth Arrest and DNA Damage-inducible 45 (GADD45) signaling. A coexpression module (thistle2) identified by weighted gene co-expression network analysis (WGCNA) was significantly correlated with alcohol dependence, alcohol consumption, and AUDIT scores. Genes in the thistle2 module were enriched with genes related to calcium signaling pathways and showed significant downregulation of these pathways, as well as enrichment for biological processes related to nicotine response and opioid signaling. A second module (brown4) showed significant upregulation of pathways related to immune signaling. Expression quantitative trait loci (eQTLs) for genes in the brown4 module were also enriched for genetic associations with alcohol dependence and alcohol consumption in large genome-wide studies included in the Psychiatric Genetic Consortium and the UK Biobank’s alcohol consumption dataset. By leveraging multi-omics data, this transcriptome analysis has identified genes and biological pathways that could provide insight for identifying therapeutic targets for alcohol dependence.

## INTRODUCTION

Alcohol dependence (AD) can be defined as a cluster of physiological, behavioral, and cognitive phenomena in which the use of alcohol takes a much higher priority for a given individual than other behaviors that once had greater value (American Psychiatric Association 1994)^1^. The development of AD is characterized by frequent episodes of intoxication, preoccupation with alcohol, use of alcohol despite adverse consequences, compulsion to seek and consume alcohol, loss of control in limiting alcohol intake, and emergence of a negative emotional state in the absence of the drug (American Psychiatric Association 1994)^1^. The changes in behavioral priorities not only results in increased morbidity and mortality, it is also a substantial social and economic burden on individual families and the nation^2^.

In individuals with alcohol dependence, there is a complex interplay between genetic background, environmental factors, and history of alcohol exposure^3^. Alcohol crosses the blood brain barrier and triggers changes in the central nervous system^4^, including transcriptional changes in many different regions of the brain ^5–9^. The transcriptional effects of long-term alcohol consumption are associated with pathways involved in the neuro-immune system, neurotoxicity, and changes in neuroplasticity ^6,7,9^. Transcriptomes from complex tissues, such as human brain, may be organized into networks of co-expressed genes that better reflect the biological functions and organization of the tissue ^7–14^. Application of bioinformatics techniques, such as weighted gene co-expression network analysis (WGCNA)^15^, has uncovered networks associated with alcohol dependence^8,9^. However, past studies were performed on small numbers of AD cases, thus limiting the statistical power to detect small changes in alcohol-induced gene expression. In this study, we utilized massively parallel sequencing of RNA transcripts from postmortem human prefrontal cortex (PFC) of 65 alcoholics and 73 controls of European descent to explore transcriptional networks and genetic variation and identified groups of coexpressed genes associated with alcohol dependence. Our analysis provides systems-level evidence of genetic networks within the PFC that contribute to the pathophysiology of alcohol drinking behavior in humans.

## MATERIALS AND METHODS

### Case selection and postmortem tissue collection

Human autopsy brain samples were obtained from the New South Wales Tissue Resource Centre at the University of Sydney (http://sydney.edu.au/medicine/pathology/btrc/). Fresh frozen samples of the superior frontal gyrus (Brodmann area 8; referred to as prefrontal cortex (PFC) in this manuscript) were collected from each postmortem sample. Brain tissue was sectioned at 3 mm intervals in the coronal plane. Alcohol dependent diagnoses were confirmed by physician interviews, review of hospital medical records, questionnaires to next-of-kin, and from pathology, radiology, and neuropsychology reports. Tissue samples were matched as closely as possible according to age, sex, post-mortem interval, pH of tissue, disease classification, and cause of death. To be included as part of the alcohol-dependent cohort, subjects had to meet the following criteria: greater than 18 years of age, no head injury at time of death, lack of developmental disorder, no recent cerebral stroke, no history of other psychiatric or neurological disorders, no history of intravenous or polydrug abuse, negative screen for AIDS and hepatitis B/C, post-mortem interval within 48 hours, and diagnosis of AD meeting the DSM-IV criteria^1^.

### Sample preparation

The Qiagen RNeasy and Lipid Tissue kit (Qiagen, Valencia, CA, USA) was used to extract total RNA from human PFC brain tissue, and RNA concentration was measured with a NanoDrop 8000 spectrophotometer (ThermoFisher Scientific). An Agilent Bioanalyzer (Agilent Technologies, Santa Clara, CA, USA) was used to test the integrity of RNA samples. Samples with an RNA integrity number (RIN) <5.5 were removed from futher analyses. Sixty samples were processed at the Waggoner Center for Alcohol and Addiction Research (WCAAR), The University of Texas at Austin while 83 samples were processed at the Ronald M. Loeb Center for Alzheimer disease, Icahn School of Medicine at Mount Sinai. Details about the library preparation and sequencing is provided in the supplementary document.

### Mapping and quantification of gene expression

Raw reads were aligned to human genome 19 (hg19) using STAR aligner (version 2.5.3.a)^16^. We used QC tools RSeQC (http://code.google.com/p/rseqc/) and Picard (https://broadinstitute.github.io/picard/) to evaluate RNA sequence quality including the %GC, %duplicates, gene body coverage, unsupervised clustering, and the library complexity. We used the Picard “MarkDuplicates” option to flag and remove duplicate reads. Gene quantification was performed with featureCounts (SUBREAD package; release 1.6.0)^17^ using Gencode annotations (Release 19 (GRCh37.p13)).

### Selection of covariates to for analyses

#### Linear regression

We first performed a linear regression with alcohol dependence as a dependent variable to identify possible covariates (e.g. sex, age, PMI). The mean age of AD subjects was 55.65 years and was not significantly different from the age of control subjects (54.96) (Table 1). There was no significant difference in distribution of RIN and brain pH between cases and controls (Table 1). Postmortem interval (PMI) was significantly lower for the alcohol dependent subjects.

**Table 1:** Demographic profile of alcohol-dependent and control subjects

| **Trait** | **Alcohol Dependent**<br>**(N = 65)** | **Control**<br>**(N = 73)** |
| --- | --- | --- |
| Male (%) | 51 (78%) | 60 (82%) |
| Mean Age (SD) (yrs) | 55.65 (11.81) | 54.96 (12.11) |
| Mean PMI (SD) (hrs) | 33.66 (15.59)* | 26.63 (13.25) |
| Brain pH (SD) | 6.54 (0.23) | 6.58 (0.29) |
| RIN (SD) | 6.84 (0.96) | 7.0 (1.01) |

\*P-value= 0.0049

**Table 2:** Results of GWAS enrichment analysis in modules correlated with alcohol dependence and alcohol consumption

|  | **RNA-Seq data (N=138)** |  |  |  |  |  | **GWAS data** |  |  |
| --- | --- | --- | --- | --- | --- | --- | --- | --- | --- |
|  | **Module trait correlation** |  |  |  |  |  | **GWAS P 0.05, eQTL P < 5 X 10<sup>-8</sup>** |  |  |
| **ID** | **AD** | **P** | **Audit** | **P** | **AC** | **P** | **PGC-AD** | **UKBB-AC** | **TAG-CPD** |
| Thistle2 | -0.28 | 9.00E-04 | -0.25 | 3.00E-03 | -0.22 | 9.00E-03 | 1.50E-02* | 1.30E-02* | 5.52E-01* |
| Brown4 | 0.18 | 4.00E-02 | 0.14 | 1.00E-01 | 0.12 | 1.00E-01 | 4.20E-03<sup>+</sup> | 2.28E-01<sup>+</sup> | 4.81E-03<sup>+</sup> |

\*Permuted P-value for the left-tail Fisher's exact test (under-enriched);
<sup>+</sup>Permuted P-value to test right-tail Fisher's exact test (over-enriched)
(AD = Alcohol Dependence; Audit = Audit scores; AC = Alcohol consumption)

#### Variance partition analysis

We used the variancePartition package^18^ in R to calculate the proportion of variance in RNA expression explained by known covariates such as age, gender, RIN and PMI, using the variancePartition package in R. The variancePartition^18^ package uses linear mixed model based statistical methods to quantify the contribution of multiple sources of variation and identify the covariates that required correction in the final analysis. Supplementary figure 1A shows violin plots depicting drivers of variation in gene expression data without accounting for covariates. The figure shows that sequencing batch is a major driver of variation in a large proportion of genes, while RIN and sex have large effects on only a few genes. We used the voom function in the Limma package (https://www.bioconductor.org/packages/devel/bioc/vignettes/limma/) to account for the effect of sequencing batch, RIN, age, sex and PMI on gene expression. After removing the effects of these covariates, alcohol-related phenotypes explained the largest proportion of the remaining variation in gene expression (Supplementry figure 1B).

### Differential gene expression analysis

Gene-level analyses started with the featureCounts-derived sample-by-gene read count matrix. The basic normalization and adjustment pipeline for the expression data matrix consisted of: (i) removal of low expression genes (< 1 CPM in > 50% of the individuals); (ii) differential gene expression analysis based upon adjustment for the chosen covariates. We filtered out all genes with lower expression in a substantial fraction of the cohort, with 18,463 genes with at least 1 CPM in at least 50% of the individuals; note that only these genes were carried forward in all subsequent analyses. The following design was used for the final differential expression analysis using the DeSeq2 ^19^ package as implimented in R: *gene expression ∼ DSM4 alcohol classification +sex + age + PMI + RIN + batch*.

### Pathway analyses of differential expression

Ingenuity® Pathway Analysis (IPA®) was used to perform pathway, canonical pathways, and causal network analysis. All genes that passed the threshold of significance at 25% FDR were included in the analysis.

### Gene ontology analysis

Gene ontology analyses were performed using the clusterProfiler package ^20^ as implemented in R. All differentially expressed genes that passed the threshold of significance at 25% FDR were included in the analysis. Results for the enrichment analysis were extracted and plotted using the ggplot2 package in R.

### Gene co-expression analysis

Scale-free co-expression networks were constructed using the weighted gene coexpression network analysis (WGCNA) package in R^15^. WGCNA provides a global perspective, emphasizing the correlation between genes to classify different molecular groupings, rather than focusing on individual genes. WGCNA defines modules using a dynamic tree-cutting algorithm based on hierarchical clustering of expression values (minimum module size=100, cutting height=0.99, deepSplit=TRUE). The networks were constructed at a soft power of 14 at which the scale free topology fit index reached 0.90 (Supplementary Figure 2B). We further merged modules that had similar co-expression patterns by calculating the eigengenes and merging those having a correlation > 75% (Supplementary Figure 2C). Correlation of module eigengenes with alcohol dependence, alcohol consumption, AUDIT scores and number of years of drinking (module-trait correlation analysis) was evaluated using Spearman’s rank correlation analysis. We used the DSM4 criteria for alcohol dependence classification as provided by the New South Wales Tissue Resource Centre at the University of Sydney. For each individual in the RNA-Seq dataset a module eigen value was calculated for each module. This module eigen value was used to perform the correlation analysis of the traits (e.g. alcohol dependence, alcohol consumption and Audit scores) with each whole module. Digital deconvolution showed no significant differences in the percentage of neurons, astrocytes and microglia in the PFC of alcoholics and controls (Supplementary Figure 3)^21^; therefore we did not perform any correction for cell-type heterogeneity. Assigned modules were functionally annotated against known molecular/functional categories and pathways using Ingenuity Pathway Analysis (IPA).

### GWAS enrichment analysis

The summary statistics from a GWAS of alcohol dependence (PGC-AD) were provided by the Psychiatric Genetics Consortium Substance Use Dependence working group^22^ (Walters et al, 2018). Summary statistics for the UKBB alcohol consumption (UKBB-AC) GWAS ^23^ were provided by Dr. Toni Clarke. We also downloaded the summary statistics for Tobacco and Genetics (TAG) Consortium’s GWAS^24^ of cigarettes per day from the PGC website (https://www.med.unc.edu/pgc/results-and-downloads). SNPs from the PGC-AD and UKBB-AC studies were mapped to PFC expression quantitative trait loci (eQTLs) in 461 post-mortem brains from the Religious Orders Study and Memory and Aging Project (ROS/MAP)^25^ (Bennett et al). Enrichment analysis was performed for SNPs meeting the criteria of eQTL P < 5 × 10^-8^ in the ROSMAP dataset and tested for overrepresentation in GWAS of AD (PGC-AD), alcohol consumption (UKBB-AC) and TAG-CPD. Since there are a few loci that passed the genome-wide significance threshold in alcohol and smoking GWAS analysis, we tested the polygenicity of alcoholism and smoking by exploring the overenrichment in variants that passed nominal threshold of significance in these datasets. The enrichment analysis was focused on eQTLs for the genes within modules that were correlated with AD in the module-trait correlation analysis.

The two modules (thistle and brown4) that showed significant enrichment (p < 0.05) in the Fisher exact test were subjected to 100,000 permutations to report the final P value of enrichment. We also performed the gene based analysis by Multi-marker Analysis of GenoMic Annotation (MAGMA)^26^ on summary statistics of PGC-AD, UKBB-AC and TAG-CPD GWAS using Functional Mapping and Annotation of GWAS (FUMA-GWAS)^27^. The summary statistics of this gene based analysis were overlaid on the IPA networks to identify the genes in these networks that also have nominal to moderate evidence of genetic contributions.

## Results

### Differential expression analysis

Analysis of PFC tissue derived from 65 alcoholics and 73 controls identified 827 differentially expressed genes at 25% FDR, 298 genes at 10% FDR and 129 genes at 5% FDR (Figure 1A, Supplemental table 1; protein coding genes only). Transient Receptor Potential Cation Channel Subfamily C Member 3 (*TRPC3*) was the top differentially expressed gene with significantly lower expression in alcohol-dependent subjects (FC 0.82; p = 4.6 x 10^-9^), while Kinesin Family Member 19 (*KFM19*) showed significantly higher expression in alcohol dependent subjects (FC 1.24; p = 5.7 x 10^-9^). IPA analysis of the differentially expressed genes (FDR < 25%) showed significant enrichment for pathways involved in interferon signaling, GADD45 signaling, and other immune-related pathways (Figure 1B). Gene-ontology enrichment analysis using clusteProfiler mapped a large proportion of genes to biological processes involved in blood coagulation and fluid transport (Figure 1C). The network analysis in IPA mapped the significant genes to networks involved in neurodegenerative disorders and organismal injury. Several genes that were part of this network were also nominally significant (p < 0.05) in the PGC-AD and UKBB-AC GWAS (Figure 1D).

**Figure 1:**
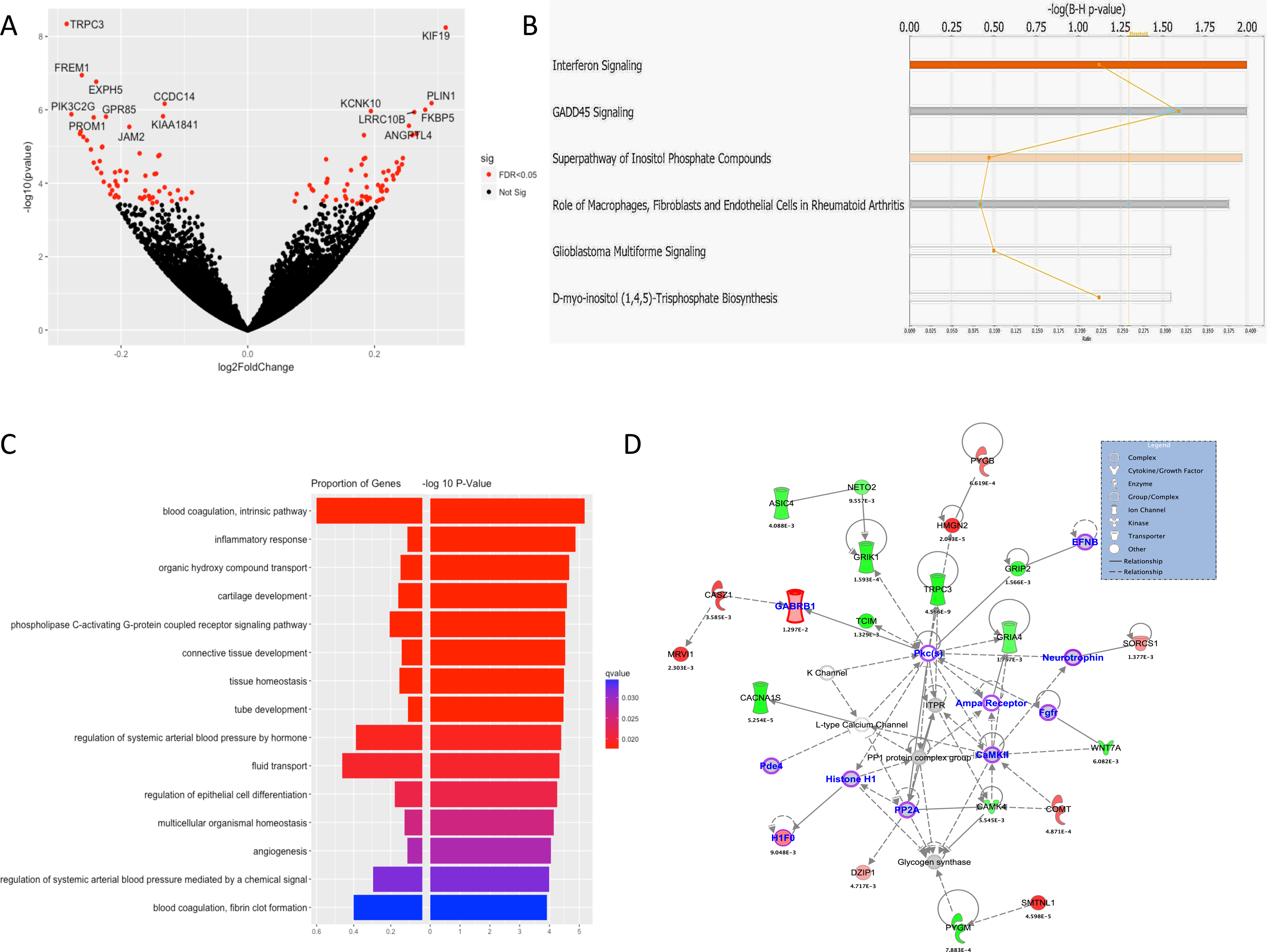
Top genes, pathways and networks from differential gene expression in DLFPC region from 68 alcoholics and 70 controls. (A) Volcano plot showing top differentially expressed genes among cases and controls. (B) The genes passing FDR threshold of 20% were inputted to IPA for pathway enrichment analysis. The figure shows some of the top pathways identified by IPA. P values here are from right tail Fisher’s exact test. (C) Enrichment analysis of gene ontology “biological process” terms. Color depicts the qvalues with red being the strongest evidence of enrichment (D) Network analysis on top genes (FDR <=20%) mapped to networks involved in the neurodegenerative disorders and organismal injuries. P value under the gene is the uncorrected p-value for differential expression among alcoholics and controls. The nominally significant genes in the UKBB-alc and PGC-SUD GWAS are highlighted with purple border and blue annotation.

### Identification of gene co-expression networks and modules

After correcting for the effects of batch, age, and RIN, the hierarchical clustering of expression data from nearly 18,000 genes generated 27 different modules (Supplementary Figure 1). Trait-module correlation analyses identified five modules that were significantly correlated to at least one alcohol related trait (Figure 2). Of these five modules, the thistle2 module (containing 72 genes), was negatively correlated with alcohol dependence and other alcohol related traits. The brown4 module (containing of 795 genes) was positively correlated with AD, AUDIT, alcohol consumption and duration of alcohol use.

**Figure 2:**
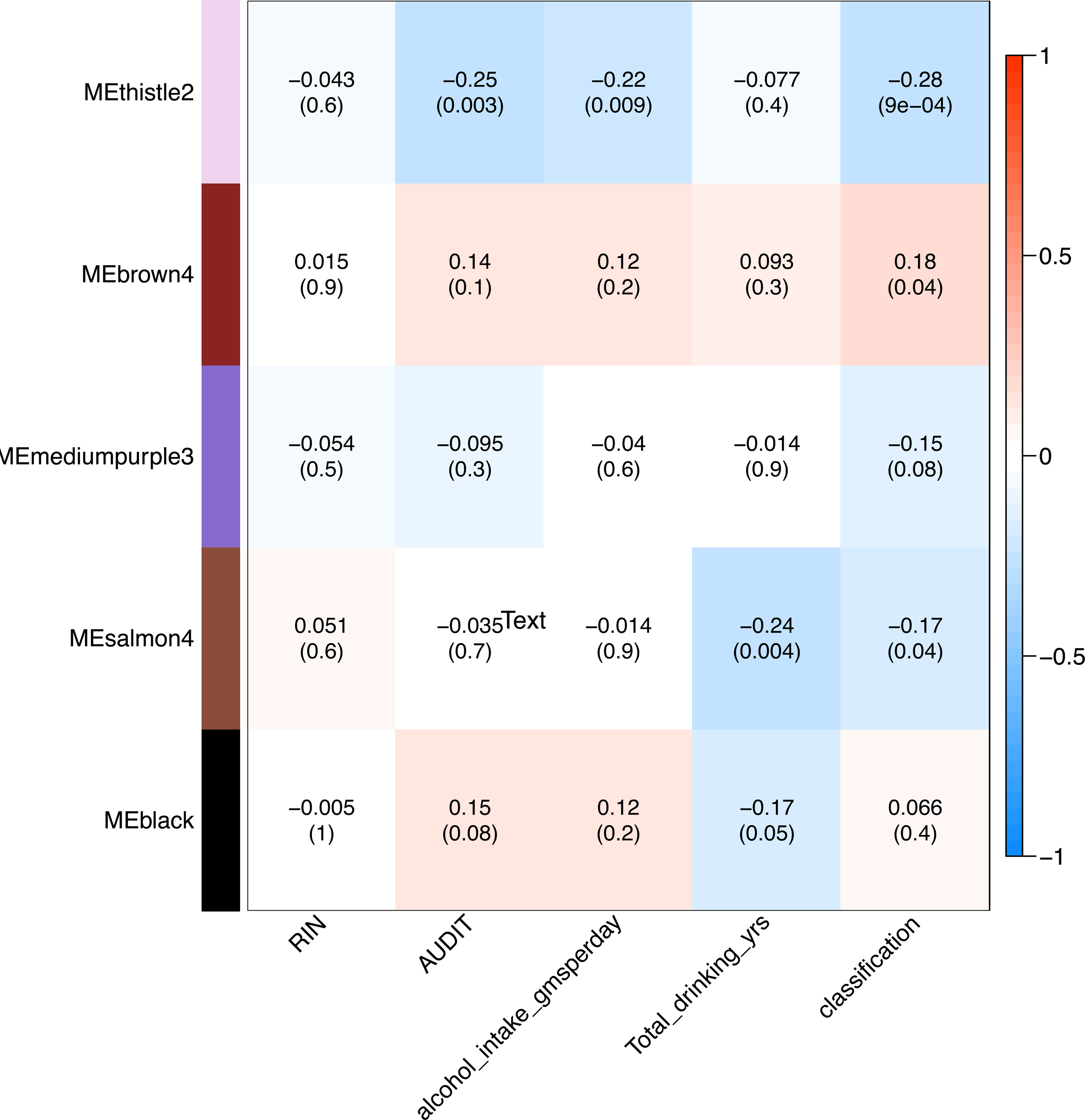
Trait module correlations with P values for the top 5 modules. WGCNA identified 27 modules, out of which 5 modules showed nominal-moderate statistical significance with any of 4 alcohol related trait (AUDIT, alcohol consumption (gms/ day), duration of drinking (years), DSM4 AD (classification). Thistle2 module also passed the multiple test correction (27 modules, 4 traits; 0.05/31 = 1.6 x 10^-3^).

#### Thistle2 module

Pathway enrichment analysis of the thistle2 module showed a significant down-regulation of pathways related to calcium signaling (Figure 3A). Gene-ontology enrichment analysis using the clusterpProfiler showed significant enrichment for biological processes involved in “response to nicotine” and “excitatory postsynaptic potential” (Figure 3B). Several genes in the thistle2 module that were significantly down-regulated in the PFC of alcohol dependent subjects. Differentially expressed genes in the thistle2 module mapped to networks involved in G-protein coupled receptor signaling, calcium signaling, and opioid signaling (Figure 3C). Cholinergic Receptor Nicotinic Alpha subunits 6 and 2 (*CHRNA6* UKBB-AC P = 7.60x 10^-3^; *CHRNA2* PGC-AD P = 1.4 x 10^-2^), Meningioma 1 (*MN1*, PGC-AD P = 9.1 x 10^-3^) and Hyaluronan And Proteoglycan Link Protein 1 (*HAPLN1*, UKBB-AC P = 1.9 x 10^-2^) are some exmples where differntialy expressed genes in thistle 2 module also showed some evidence of genetic contribution towards alcohol consumption or dependence.

**Figure 3:**
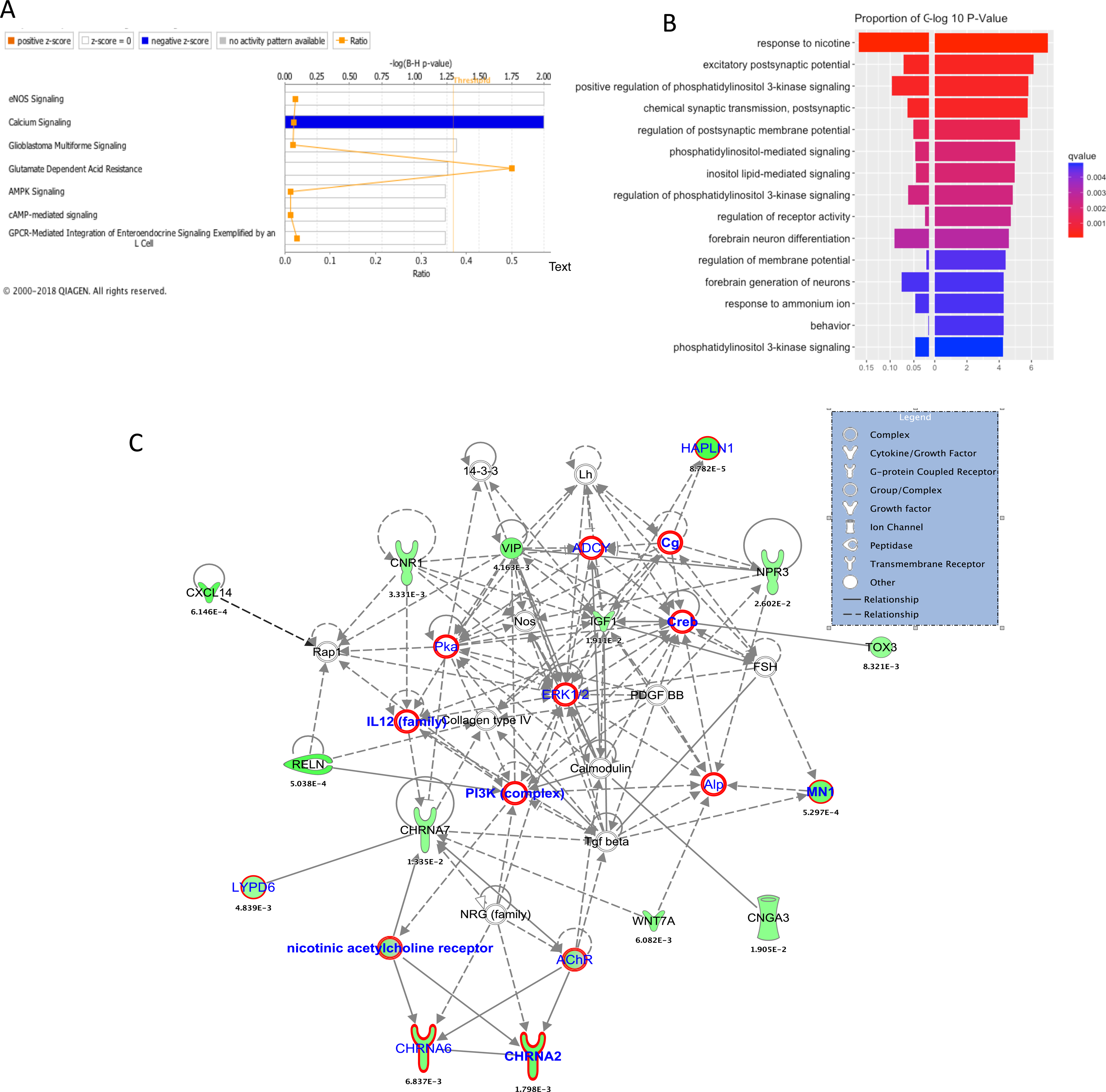
Enrichment analysis of genes in thistle2 module that are differentially expressed in alcoholics and controls. (A) More than 50% of genes in calcium signaling pathways were found to be down-regulated in the thistle2 module. (B) Enrichment analysis for GO:BP terms showed downregulation of genes related to response to nicotine and postsynaptic potential. (C) Nearly 15 genes mapped to network related to amino-acid metabolism with many genes that were involved in G-protein coupled receptor signalling, calcium signaling and opioid signaling pathway. The nominally significant genes in the UKBB-alc and PGC-SUD GWAS are marked with red boundaries (*ADCY5* P = 7.07 x 10^-7^ in UKBB-AC, *ADCY7*, P = 2.2 x 10^-4^ in UKBB-AC), *IL12B*, P = 1.1 x 10^-2^ in PGC-AD, *PIK3C2G*, P = 6.8 x 10^-3^ in UKBB-AC, *PIK3R4*, P = 3.4 x 10^-2^ in PGC-AD, *CHRNA6* in UKBB-AC P = 7.60x 10^-3^, *CHRNA2* in PGC-AD P = 1.4 x 10^-2^, *MN1* in PGC-AD P = 9.1 x 10^-3^ and *HAPLN1* in UKBB-AC P = 1.9 x 10^-2^).

#### Brown4 module

Pathway analysis for differentially expressed genes in the brown4 module showed significant enrichment for Growth Arrest and DNA Damage (GADD45) signaling and for biological processes related to the inflammatory response (Figure 4). Other genes that were also significantly upregulated in the PFC of alcoholics mapped to networks involved in infectious and respiratory diseases.

**Figure 4:**
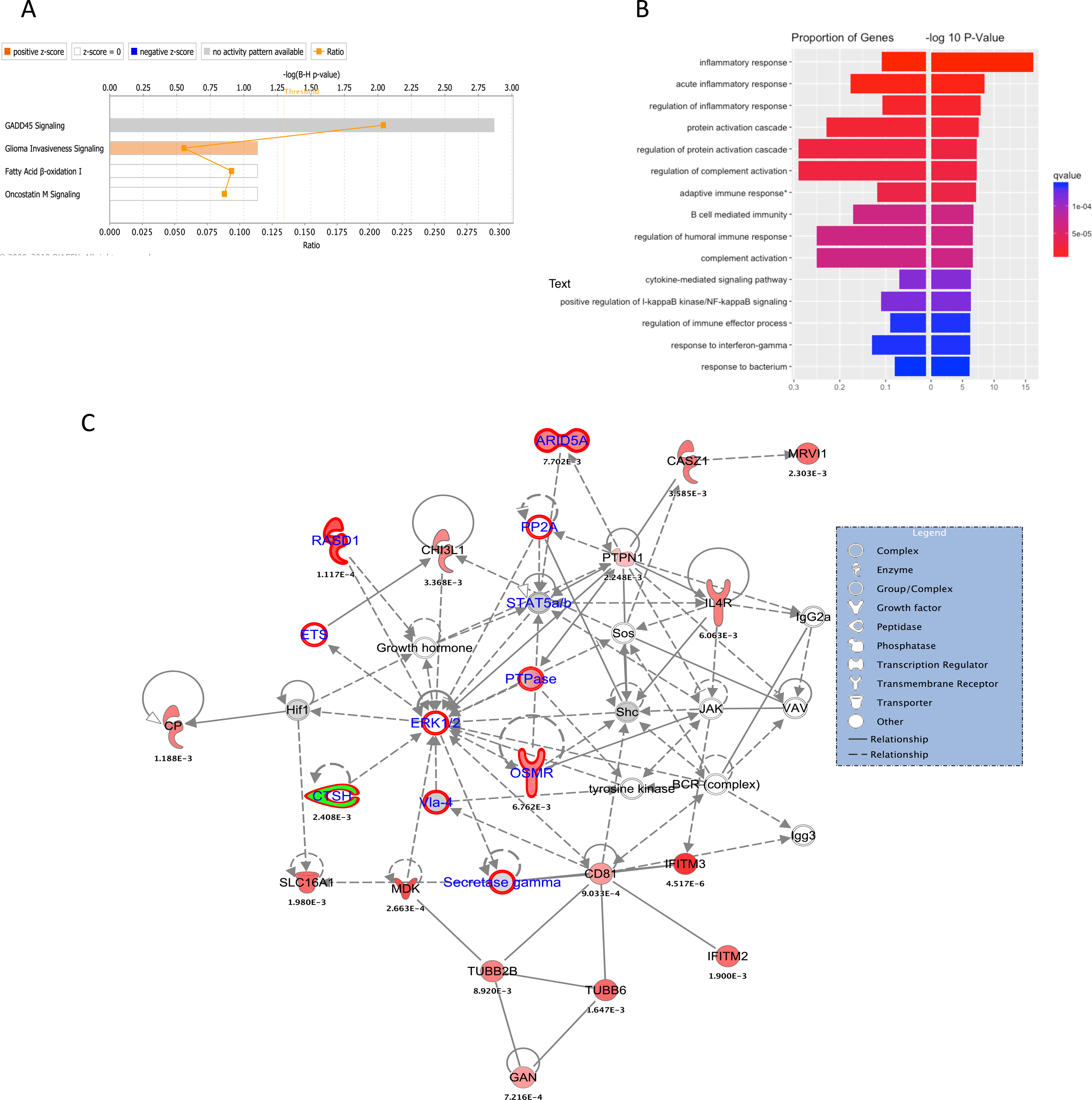
Enrichment analysis of brown4 module genes that were differentially expressed genes (FDR* < 0.05) among alcoholics and controls. (A) Pathway analysis showed significant upregulation of genes related immune signaling and metabolism. (B) Enrichment analysis for GO:BP terms showed enrichment of genes related to inflamatory response. (C) The genes in the brown4 module mapped to network involved in infectious and respiratory diseases. The genes that were nominally significant in the UKBB-Alc and PGC-SUD GWAS are highlighted with purple.

### GWAS enrichment analysis

GWAS enrichment analysis of significant eQTLs (P < 5 × 10^-8^) for all genes in the top 5 modules (ranked by P value in module-trait correlation analysis), showed evidence of enrichment for SNPs associated with AD (GWAS p < 0.05) in PGC-AD and alcohol consumption in UKBB-AC datasets. The brown4 module was also enriched for GWAS association in the TAG-CPD dataset. The thistle2 module did not show enrichment of GWAS association. Surprisingly, genes in the thistle2 modules were significantly depleted for GWAS signals in the PGC-AD and UKBB-AC GWAS analyses. This finding was confirmed by permutation analysis.

### Discussion

To our knowledge, this is the largest transcriptome analysis comparing PFC of alcohol-dependent cases and controls. The present study identified 129 genes (FDR < 0.05) that were differentially expressed in alcohol dependent subjects (Supplementary table 1). *FKBP5*, a well studied gene that is asoociated with alcohol use^28–31^, showed increased expression in the PFC of alcohol dependent subjects in our differential gene expression analysis (l_2_FC 0.27; P = 4.57 x 10^-7^). Other studies have also shown that *FKBP5* plays a role in alcohol drinking behaviors in rodents^28,29^ and humans^32^. *FKBP5* encodes FK506-binding protein 5, a glucocorticoid receptor (GR)-binding protein implicated in various psychiatric disorders and alcohol withdrawal severity^30^. Qiu and colleagues^30^ reported that *Fkbp5* KO mice exhibited increased alcohol consumption compared with wild-type mice. Another study has shown that the absence of *Fkbp5* enhances sensitivity to alcohol withdrawal in mice^33^. Recent findings also suggested that *Fkbp5* expression in mesocorticolimbic dopaminergic regions is associated with early life-stress mediated sensitivity to alcohol drinking and that there is a gene environment interaction among *FKBP5* genotype and parent-child relationship that influences alcohol drinking.

Genes showing significant differences in expression between alcohol dependent subjects and controls were enriched in pathways related to interferon and GADD45 signaling (Figure 1 B). Interferons are cytokines that have antiviral, antiproliferative, and immunomodulatory effects and the interferon pathway plays a critical role in human innate and adaptive immune responses^34^. Our pathway analysis results are consistent with earlier findings showing induction of innate immune genes by stress and drug abuse ^35,35^. Furthermore, mRNA expression studies in human brain showed significant changes in expression of genes related to immune or inflammatory responses in hippocampus^7^ and nucleus accumbens^8^. The neuroinflammation associated with chronic alcohol exposure and withdrawal may be attributed to microglial activation and is associated with the neuropathology of chronic alcohol exposure^36^. Differentially expressed genes (FDR < 25%) also mapped to networks associated with neurodegenerative disorders and organismal injury (Figure 1 D). Many differentially expressed genes in this network are involved in nervous system development and function. Specifically *TRPC3* and calcium dependent protein kinase 4 (*CAMK4*) are involved in excitatory post-synaptic current while Ampa receptor, Glutamate Ionotropic receptor AMPA type subunit 4 (*GRIA4*), Calcium dependent protein kinase ii (*CaMKII*) and *CAMK4* are involved in synaptic transmission.

Although we identified several genes that were differentially expressed in the PFC of alcohol-dependent subjects, the variance explained by individual genes was very small (0.15-1%). The differential expression observed here is smaller than that reported in earlier differential expression studies of alcoholism, but it is consistent with differential expression studies of larger sample size^37^. For example, the CommonMind consortium reported similar fold changes in the differential expression study of schizophrenia and they showed that their observation is consistent with plausible models for average differential gene expression and the polygenic inheritance of schizophrenia. The polygenicity of AD has also been observed by the GWAS of alcoholism and other complex behavioral/psychiatric disorders ^22,38–41^, and it was demonstrated that effect size for each individual genetic variant is very small. Studies that used a co-expression network approach also showed that alcohol dependence is shaped, in part, by persistent alterations in networks of co-expressed genes that collectively mediate excessive drinking and other alcohol-dependent phenotypes^8,9^. These and other studies also demonstrated that the gene network structure is significantly correlated with lifetime alcohol consumption in addition to an overall loss in network structure; furthermore, the neurobiology of alcohol dependence may be due to altered covariation of gene modules, rather than discrete changes in differentially expressed genes across the transcriptome^9,13^.

Trait-module correlation analysis for the thistle2 module showed a significant negative correlation with alcohol dependence (-0.28, P = 9.0 x 10^-4^), alcohol consumption (-0.22, P = 9.0 x 10^-3^), and AUDIT score (-0.25, P = 3.0 x 10^-3^), while the brown4 module showed a positive correlation (0.18, P = 4.0 x 10^-2^) with alcohol dependence (Figure 2). The salmon4 module was associated with the total number of drinking years (-0.24, P = 4.0 x 10^-3^), independent of the age of the subjects. Genes in the thistle2 module were significantly down-regulated in the PFC from alcoholics. Many genes in the thistle2 module mapped to networks involved in opioid signaling and nicotine response, highlighting the importance of this module in addiction-related traits.

Pathway analysis showed that all genes that overlapped with genes involved in calcium signaling were significantly downregulated (Figure 3A). Acute ethanol exposure has been shown to inhibit Ca^2+^ currents induced by PKC-dependent phosphorylation of mGluR5 in neurons^42^. Early studies in PC12 cell cultures also showed that ethanol has a significant inhibitory effect on the influx of Ca^2+^ through L-type voltage-gated Ca^2+^ channels^43^. Alcohol exposure also modulates Ca^2+^ signaling between astrocytes and neurons^44^ (Warden et al, 2016), and Ca^2+^ acts as a second messenger that controls multiple processes in immune cells, including chemotaxis and secretion of pro- and anti-inflammatory cytokines. Our analyses provide further evidence that alcohol exposure alters Ca^2+^ signaling in the brains of alcoholics and could potentially alter communication between neurons and brain immune cells. Another module that correlated with alcohol dependence, brown4, was also enriched in immune response and infectious diseases, providing additional evidence for the role of the neuroimmune system in the etiology of alcohol dependence. Some of the differentially expressed genes in this network were also statistical significant in the gene-based tests (*RASD1*, UKBB-AC, P = 1.64 x 10^-5^ and *ARID5A*, UKBB-AC, P 1.4 x 10^-3^). The differentially expressed *FKBP5* gene was also part of the brown module, but it was not identified as hub gene according to intra-modular connectivity (supplementary table 2).

Enrichment analysis of nominally significant GWAS variants (p < 0.05) that were also eQTLs (p < 5 × 10^-8^) for genes in the thistle2 module showed significant under enrichment in the two-tail Fisher test. The under-enrichment remained significant even after 100,000 permutations. This might be due to the small size of this module (N = 72 genes). Although some of the differntialy expressed genes were significant in the gene-based tests performed in UKBB-AC and PGC-AD datasets using MAGMA (*CHRNA6, CHRNA2, MN1* and *HAPLN1*). In the calcium signaling network (Figure 3 C), a few genes that were not part of the thistle2 module, but were essential to create network connections, were also found to be significant (3.4 x 10^-2^ < P > 4.8 x 10^-2^) in the gene-based tests (circled in red; Fig 4C). This suggests possible gene-environment (alcohol exposure) interactions in the etiology of alcohol dependence. This also reinforces the need for multi-omics data to understand a complex disorder like alcoholism. eQTLs for genes in the brown4 module (N = 726 genes) were significantly enriched for GWAS signals (P = 4.2 x 10^-3^) in the PGC-AD GWAS. Interestingly this module was also positively correlated with alcohol dependence (0.18, P = 4.0 x 10^-2^) in trait-module correlation analysis.

Because of limited availability of human post-mortem tissue with DSM4 alcohol dependence phenotype, we tried to look for validation in rodent RNA-expression datasets (Supplementary methods; supplementary table 2). The hub genes identified in present analysis were found to be significant enriched for association signals in the rodents. This observation adds to the validity of hub genes in the identified modules.

In the present study, we focused on integrating the genomic information to transcriptomic data to identify gene (genetic background) x environment (alcohol exposure) interactions in the etiology of alcohol use disorders. As mentioned in the discussion we identified that genes that have altered expression due to alcohol exposure interact with risk genes (GWAS) to increase an individual’s risk of becoming dependent on alcohol. So, to translate these findings in animals, one has to mimic expression of hub genes as well as the risk gene to alter the pathways associated with alcoholism. We are also reporting the direction of effect of the differential expression. That should provide information that can be used to see whether knock-down or overexpression of key genes alters risk for AUD phenotypes in models. Also, the replication of the modules in rodent models indicates which models might be useful to study the effects of dysregulation in these models.

Multiple lines of evidence derived from this study allowed us to prioritize the genes altered by exposure to alcohol. The gene co-expression network analysis also identified networks of genes altered in alcohol-dependent subjects. Further support for our findings comes from work showing that many genes in these networks were also associated with alcohol dependence and alcohol consumption in large GWAS study cohorts. This systematic exploration of transcriptomic organization in the PFC from alcoholics provides further support for the role of the neuroimmune system in alcohol dependence. The biological pathways and networks of genes identified in the current study will help prioritize genes for functional studies and may help advance targeted treatment approaches for alcohol use disorders.

## Conflict of interest

Dr. Alison Goate and Dr. Jen-Chyong Wang are listed as inventors on Issued U.S. Patent 8080,371, “Markers for Addiction” covering the use of certain variants in determining the diagnosis, prognosis, and treatment of addiction. Other authors report no conflict of interest.

## Funding and Disclosure

COGA is supported by NIH Grant U10AA008401 from the National Institute on Alcohol Abuse and Alcoholism (NIAAA) and the National Institute on Drug Abuse (NIDA).

Mayfield grant support: U01 AA020926 and R01 AA012404

The work was also supported in part by the Integrative Neuroscience Initiative on Alcoholism (INIA)-Neuroimmue consortium (5U24AA025479-02).

## Acknowledgments

COGA: The Collaborative Study on the Genetics of Alcoholism (COGA), Principal Investigators B. Porjesz, V. Hesselbrock, H. Edenberg, L. Bierut, includes eleven different centers: University of Connecticut (V. Hesselbrock); Indiana University (H.J. Edenberg, J. Nurnberger Jr., T. Foroud; Y. Liu); University of Iowa (S. Kuperman, J. Kramer); SUNY Downstate (B. Porjesz); Washington University in St. Louis (L. Bierut, J. Rice, K. Bucholz, A. Agrawal); University of California at San Diego (M. Schuckit); Rutgers University (J. Tischfield, A. Brooks); Department of Biomedical and Health Informatics, The Children’s Hospital of Philadelphia; Department of Genetics, Perelman School of Medicine, University of Pennsylvania, Philadelphia PA (L. Almasy), Virginia Commonwealth University (D. Dick), Icahn School of Medicine at Mount Sinai (A. Goate), and Howard University (R. Taylor). Other COGA collaborators include: L. Bauer (University of Connecticut); J. McClintick, L. Wetherill, X. Xuei, D. Lai, S. O’Connor, M. Plawecki, S. Lourens (Indiana University); G. Chan (University of Iowa; University of Connecticut); J. Meyers, D. Chorlian, C. Kamarajan, A. Pandey, J. Zhang (SUNY Downstate); J.-C. Wang, M. Kapoor, S. Bertelsen (Icahn School of Medicine at Mount Sinai); A. Anokhin, V. McCutcheon, S. Saccone (Washington University); J. Salvatore, F. Aliev, B. Cho (Virginia Commonwealth University); and Mark Kos (University of Texas Rio Grande Valley). A. Parsian and H. Chen are the NIAAA Staff Collaborators.

We continue to be inspired by our memories of Henri Begleiter and Theodore Reich, founding PI and Co-PI of COGA, and also owe a debt of gratitude to other past organizers of COGA, including Ting-Kai Li, P. Michael Conneally, Raymond Crowe, and Wendy Reich, for their critical contributions. This national collaborative study is supported by NIH Grant U10AA008401 from the National Institute on Alcohol Abuse and Alcoholism (NIAAA) and the National Institute on Drug Abuse (NIDA).

The authors are also grateful to the New South Wales Tissue Resource Centre at the University of Sydney for providing human brain samples; the Centre is supported by the National Health and Medical Research Council of Australia, Schizophrenia Research Institute, and National Institute on Alcohol Abuse and Alcoholism (NIH/NIAAA R24AA012725).

## Data sharing policy

RNA-Seq data for alcohol dependence cases and controls will be available through GEO. All the summary statistics for differential expression will also be posted at INIA and COGA’s homepage (and Shiny web app) and will be freely available to download after publication is online.

Shiny webapp link for entire summary statistics: https://lcad.shinyapps.io/coga-inia/

## References

1. Bell CC. DSM-IV: Diagnostic and Statistical Manual of Mental Disorders. JAMA 1994; 272: 828–829.

2. Sacks JJ, Gonzales KR, Bouchery EE, Tomedi LE, Brewer RD. 2010 National and State Costs of Excessive Alcohol Consumption. Am J Prev Med 2015; 49: e73–e79.

3. Contet C. Gene Expression Under the Influence: Transcriptional Profiling of Ethanol in the Brain. Curr Psychopharmacol 2012; 1: 301–314.

4. Alfonso-Loeches S, Guerri C. Molecular and behavioral aspects of the actions of alcohol on the adult and developing brain. Crit Rev Clin Lab Sci 2011; 48: 19–47.

5. Fan L, van der Brug M, Chen W, Dodd PR, Matsumoto I, Niwa S et al. Increased expression of mitochondrial genes in human alcoholic brain revealed by differential display. Alcohol Clin Exp Res 1999; 23: 408–413.

6. Mayfield RD, Harris RA, Schuckit MA. Genetic factors influencing alcohol dependence. Br J Pharmacol 2008; 154: 275–287.

7. McClintick JN, Xuei X, Tischfield JA, Goate A, Foroud T, Wetherill L et al. Stress-response pathways are altered in the hippocampus of chronic alcoholics. Alcohol Fayettev N 2013; 47: 505–515.

8. Mamdani M, Williamson V, McMichael GO, Blevins T, Aliev F, Adkins A et al. Integrating mRNA and miRNA Weighted Gene Co-Expression Networks with eQTLs in the Nucleus Accumbens of Subjects with Alcohol Dependence. PloS One 2015; 10: e0137671.

9. Farris SP, Arasappan D, Hunicke-Smith S, Harris RA, Mayfield RD. Transcriptome organization for chronic alcohol abuse in human brain. Mol Psychiatry 2015; 20: 1438–1447.

10. Oldham MC, Konopka G, Iwamoto K, Langfelder P, Kato T, Horvath S et al. Functional organization of the transcriptome in human brain. Nat Neurosci 2008; 11: 1271–1282.

11. Flatscher-Bader T, Wilce PA. Chronic smoking and alcoholism change expression of selective genes in the human prefrontal cortex. Alcohol Clin Exp Res 2006; 30: 908–915.

12. Mayfield J, Ferguson L, Harris RA. Neuroimmune signaling: a key component of alcohol abuse. Curr Opin Neurobiol 2013; 23: 513–520.

13. Farris SP, Mayfield RD. RNA-Seq reveals novel transcriptional reorganization in human alcoholic brain. Int Rev Neurobiol 2014; 116: 275–300.

14. Hermann D, Hirth N, Reimold M, Batra A, Smolka MN, Hoffmann S et al. Low µ-Opioid Receptor Status in Alcohol Dependence Identified by Combined Positron Emission Tomography and Post-Mortem Brain Analysis. Neuropsychopharmacol Off Publ Am Coll Neuropsychopharmacol 2017; 42: 606–614.

15. Langfelder P, Horvath S. WGCNA: an R package for weighted correlation network analysis. BMC Bioinformatics 2008; 9: 559.

16. Dobin A, Davis CA, Schlesinger F, Drenkow J, Zaleski C, Jha S et al. STAR: ultrafast universal RNA-seq aligner. Bioinformatics 2013; 29: 15–21.

17. Liao Y, Smyth GK, Shi W. featureCounts: an efficient general purpose program for assigning sequence reads to genomic features. Bioinforma Oxf Engl 2014; 30: 923–930.

18. Hoffman GE, Schadt EE. variancePartition: interpreting drivers of variation in complex gene expression studies. BMC Bioinformatics 2016; 17: 483.

19. Love MI, Huber W, Anders S. Moderated estimation of fold change and dispersion for RNA-seq data with DESeq2. Genome Biol 2014; 15. doi:10.1186/s13059-014-0550-8.

20. Yu G, Wang L-G, Han Y, He Q-Y. clusterProfiler: an R Package for Comparing Biological Themes Among Gene Clusters. OMICS J Integr Biol 2012; 16: 284–287.

21. Li Z, Del-Aguila JL, Dube U, Budde J, Martinez R, Black K et al. Genetic variants associated with Alzheimer’s disease confer different cerebral cortex cell-type population structure. Genome Med 2018; 10: 43.

22. Walters RK, Adams MJ, Adkins AE, Aliev F, Bacanu S-A, Batzler A et al. Trans-ancestral GWAS of alcohol dependence reveals common genetic underpinnings with psychiatric disorders. bioRxiv 2018;: 257311.

23. Clarke T-K, Adams MJ, Davies G, Howard DM, Hall LS, Padmanabhan S et al. Genome-wide association study of alcohol consumption and genetic overlap with other health-related traits in UK Biobank (N=112?117). Mol Psychiatry 2017; 22: 1376–1384.

24. Tobacco and Genetics Consortium. Genome-wide meta-analyses identify multiple loci associated with smoking behavior. Nat Genet 2010; 42: 441–447.

25. Bennett DA, Schneider JA, Buchman AS, Barnes LL, Boyle PA, Wilson RS. Overview and findings from the rush Memory and Aging Project. Curr Alzheimer Res 2012; 9: 646–663.

26. de Leeuw CA, Mooij JM, Heskes T, Posthuma D. MAGMA: Generalized Gene-Set Analysis of GWAS Data. PLoS Comput Biol 2015; 11. doi:10.1371/journal.pcbi.1004219.

27. Watanabe K, Taskesen E, Bochoven A van, Posthuma D. Functional mapping and annotation of genetic associations with FUMA. Nat Commun 2017; 8: 1826.

28. Kerns RT, Ravindranathan A, Hassan S, Cage MP, York T, Sikela JM et al. Ethanol-Responsive Brain Region Expression Networks: Implications for Behavioral Responses to Acute Ethanol in DBA/2J versus C57BL/6J Mice. J Neurosci 2005; 25: 2255–2266.

29. Treadwell JA, Singh SM. Microarray Analysis of Mouse Brain Gene Expression Following Acute Ethanol Treatment. Neurochem Res 2004; 29: 357–369.

30. Qiu B, Luczak SE, Wall TL, Kirchhoff AM, Xu Y, Eng MY et al. The FKBP5 Gene Affects Alcohol Drinking in Knockout Mice and Is Implicated in Alcohol Drinking in Humans. Int J Mol Sci 2016; 17. doi:10.3390/ijms17081271.

31. Nylander I, Todkar A, Granholm L, Vrettou M, Bendre M, Boon W et al. Evidence for a Link Between Fkbp5/FKBP5, Early Life Social Relations and Alcohol Drinking in Young Adult Rats and Humans. Mol Neurobiol 2017; 54: 6225–6234.

32. Xie P, Kranzler HR, Poling J, Stein MB, Anton RF, Farrer LA et al. Interaction of FKBP5 with childhood adversity on risk for post-traumatic stress disorder. Neuropsychopharmacol Off Publ Am Coll Neuropsychopharmacol 2010; 35: 1684–1692.

33. Huang M-C, Schwandt ML, Chester JA, Kirchhoff AM, Kao C-F, Liang T et al. FKBP5 Moderates Alcohol Withdrawal Severity: Human Genetic Association and Functional Validation in Knockout Mice. Neuropsychopharmacology 2014; 39: 2029–2038.

34. Platanias LC. Mechanisms of type-I-and type-II-interferon-mediated signalling. Nat Rev Immunol 2005; 5: 375–386.

35. Crews FT, Zou J, Qin L. Induction of innate immune genes in brain create the neurobiology of addiction. Brain Behav Immun 2011; 25 Suppl 1: S4–S12.

36. Kalinin S, González-Prieto M, Scheiblich H, Lisi L, Kusumo H, Heneka MT et al. Transcriptome analysis of alcohol-treated microglia reveals downregulation of beta amyloid phagocytosis. J Neuroinflammation 2018; 15: 141.

37. Fromer M, Roussos P, Sieberts SK, Johnson JS, Kavanagh DH, Perumal TM et al. Gene Expression Elucidates Functional Impact of Polygenic Risk for Schizophrenia. Nat Neurosci 2016; 19: 1442–1453.

38. Agrawal A, Bierut LJ. Identifying genetic variation for alcohol dependence. Alcohol Res Curr Rev 2012; 34: 274–281.

39. Edenberg HJ, Foroud T. The genetics of alcoholism: identifying specific genes through family studies. Addict Biol 2006; 11: 386–396.

40. Gelernter J, Kranzler HR, Sherva R, Almasy L, Koesterer R, Smith AH et al. Genome-wide association study of alcohol dependence:significant findings in African- and European-Americans including novel risk loci. Mol Psychiatry 2014; 19: 41–49.

41. Kapoor M, Chou Y-L, Edenberg HJ, Foroud T, Martin NG, Madden P a. F et al. Genome-wide polygenic scores for age at onset of alcohol dependence and association with alcohol-related measures. Transl Psychiatry 2016; 6: e761.

42. Netzeband JG, Gruol DL. Modulatory effects of acute ethanol on metabotropic glutamate responses in cultured Purkinje neurons. Brain Res 1995; 688: 105–113.

43. Mullikin-Kilpatrick D, Mehta ND, Hildebrandt JD, Treistman SN. Gi is involved in ethanol inhibition of L-type calcium channels in undifferentiated but not differentiated PC-12 cells. Mol Pharmacol 1995; 47: 997–1005.

44. Warden A, Erickson E, Robinson G, Harris RA, Mayfield RD. The neuroimmune transcriptome and alcohol dependence: potential for targeted therapies. Pharmacogenomics 2016; 17: 2081–2096.

