## Supplement material for "Analysis of whole genome-transcriptomic organization in brain to identify genes associated with alcoholism"

### Supplementary Methods:

#### Sequencing

#### Total RNA sequencing at the Waggoner Center for Alcohol and Addiction Research

Total RNA was extracted from frozen PFC the kit using mirVana™ miRNA Isolation Kit, with phenol (#AM1560, Thermofisher Scientific, Waltham, MA). RNA samples were DNAse treated with DNA free kit (#AM1906, Thermofisher Scientific, Waltham, MA) and ribosomal RNA was depleted using RiboMinus Eukaryote kit (Life Technologies, Foster City, CA, USA). Sixty samples from 30 alcoholics and 30 controls were processed using the TruSeq RNA Library Prep Kit v2 and sequenced on the Illumina HiSeq 2000 at the Genome Sequencing and Analysis Facility at The University of Texas at Austin. Paired-end libraries with an average insert size of 180 bp were obtained.

#### Total RNA sequencing at the New York Genome Centre

DNase-treated RNA from 83 frozen PFC samples were provided to the New York Genome Center (NYGC) for total RNA sequencing. To eliminate preparation batch effects, all RNA samples were prepared in a single batch by one operator using the KAPA Stranded RNA-Seq Kit with RiboErase. The library quality control (QC) included a measurement of the average size of library fragments using the FragmentAnalyzer and estimation of the total concentration of DNA by PicoGreen. Paired end sequencing was performed on the Illumina HiSeq 2500 instrument. Assessment of the quality of the sequencing data included multiple metrics at several steps of the analysis pipeline. Following the completion of a sequencing run, a QC specialist reviewed the following sequencing quality metrics: number of pass filter reads per sample, base quality per cycle, percent base content per cycle, and the overall distribution of base quality scores. One sample was an intentional overlap with a sample processed at the Waggoner Center for Alcohol and Addiction Research. The duplicate samples clustered together in unsupervised clustering, and we kept the NYGC sample for further analysis. We removed four brain samples from patients that died from infections. Final analysis was performed on 138 brains (65 from alcoholics and 73 from control subjects).

**Hub Gene identification:**

In the study, hub genes were defined by module connectivity, measured by absolute value of the Pearson's correlation (ModuleMembership > 0.8). We identified hub genes in the thistle 2 and brown 4 modules that were highly correlated with certain clinical trait.

**SUPPLEMENTARY Figures (in PDF format)**

**Supplementary figure 1: Violin plots depicting drivers of variation in gene expression.** X-axis shows various traits/ confounders and y-axis shows the percentage of variance explained. Thickness of each violin bar depicts the proportion of genes corresponding to level of y-axis. (A) Sequencing batch was the major driver of variation in gene expression. (B) The variation partition plot after correction for batch, RIN, age, gender and PMI.

**Supplementary Figure 2: WGCNA derived modules of gene expression.** (A) Summary network indices (y-axes) as functions of the soft thresholding power (x-axes). Numbers in the plots indicate the corresponding soft thresholding powers. The plots indicate that approximate scale-free topology is attained around the soft-thresholding power of 14. (B) Gene dendrogram obtained by clustering the dissimilarity based on consensus Topological Overlap. The two color rows show the preliminary (unmerged) and the final, merged module assignments. (C) Correlation among module eigen values and alcohol related traits. Values in the parenthesis are the p values associated with correlation coefficient.

**Supplementary Figure 3: Deconvolution of cell types in alcoholics and controls.**

**Supplementary Figure 4: Relationship among the network connectivity and differential expression**

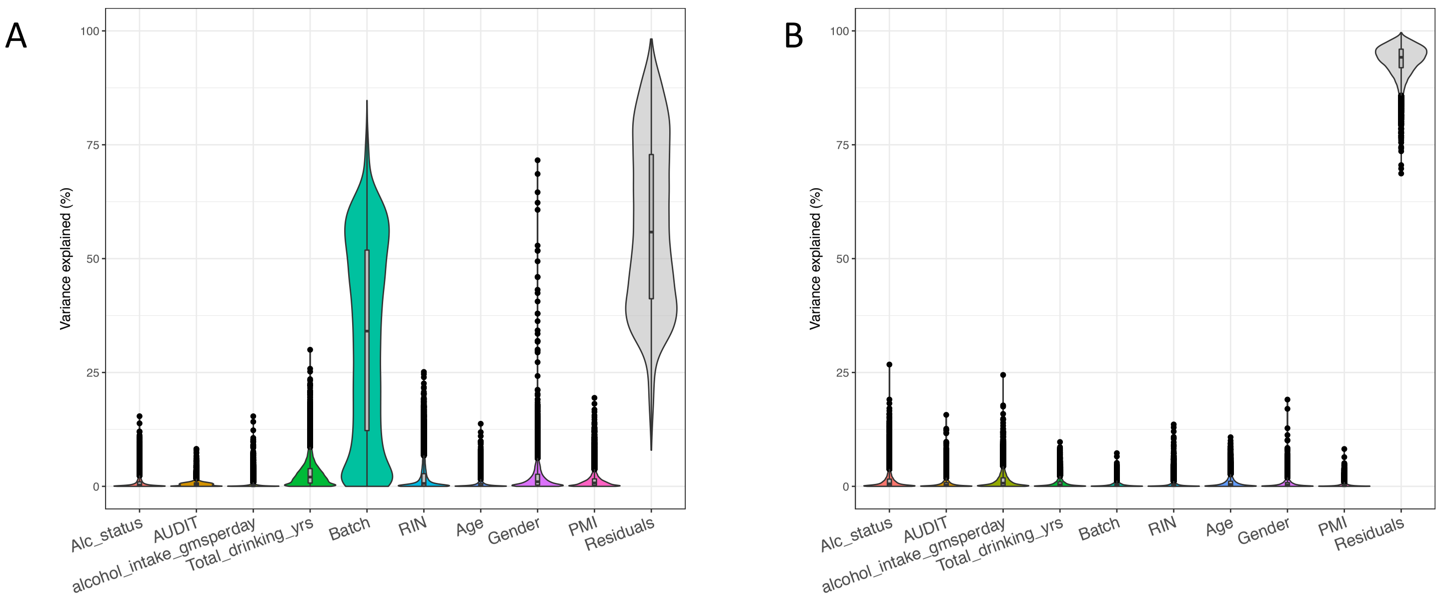

Supplementary figure 1: Violin plots depicting drivers of variation in gene expression. X-axis shows various traits/ confounders and y-axis shows the percentage of variance explained. Thickness of each violin bar depicts the proportion of genes corresponding to level of y-axis. (A) Sequencing batch was the major driver of variation in gene expression. (B) The variation partition plot after correction for batch, RIN, age, gender and PMI.

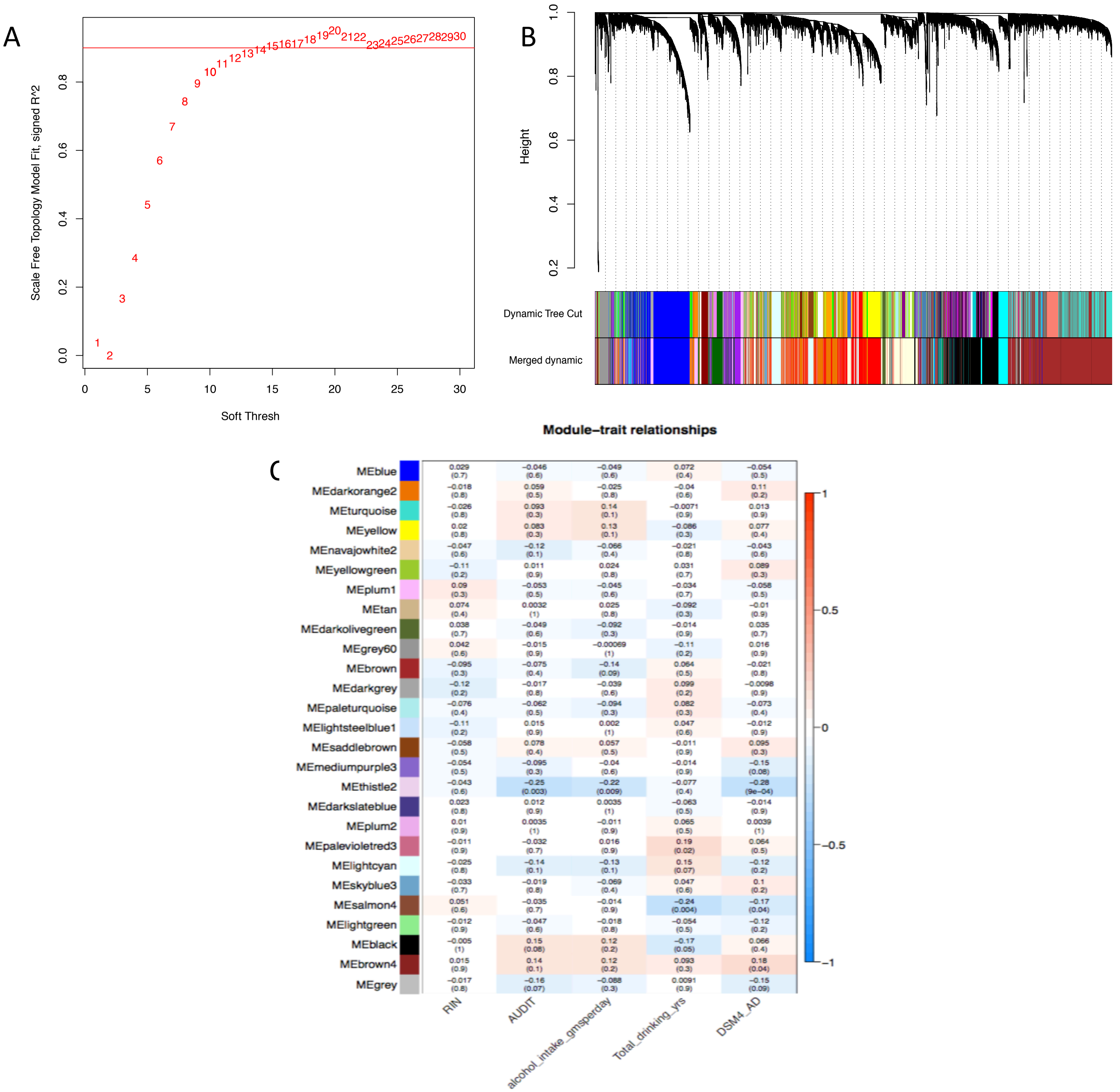

Supplementary Figure 2: **WGCNA derived modules of gene expression**. (A) Summary network indices (y-axes) as functions of the soft thresholding power (x-axes). Numbers in the plots indicate the corresponding soft thresholding powers. The plots indicate that approximate scale-free topology is attained around the soft-thresholding power of 14. (B) Gene dendrogram obtained by clustering the dissimilarity based on consensus Topological Overlap. The two color rows show the preliminary (unmerged) and the final, merged module assignments. (C) Correlation among module eigen values and alcohol related traits. Values in the parenthesis are the p values associated with correlation coeficient.

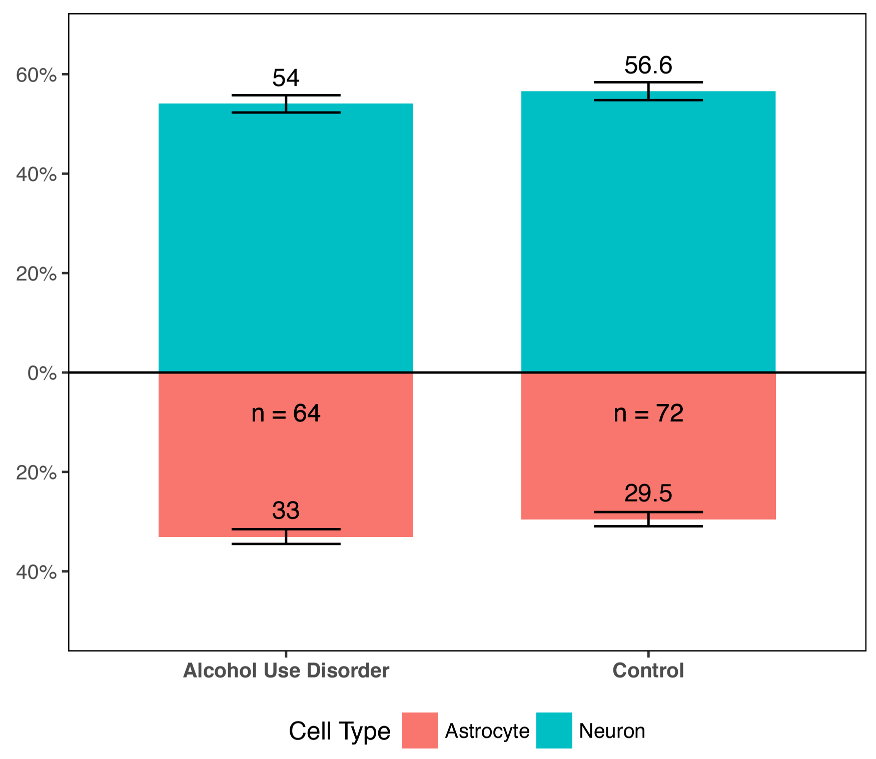

Supplementary Figure 3: Deconvolution of cell types in alcoholics and controls.

Supplementary
Figure 4 (a)

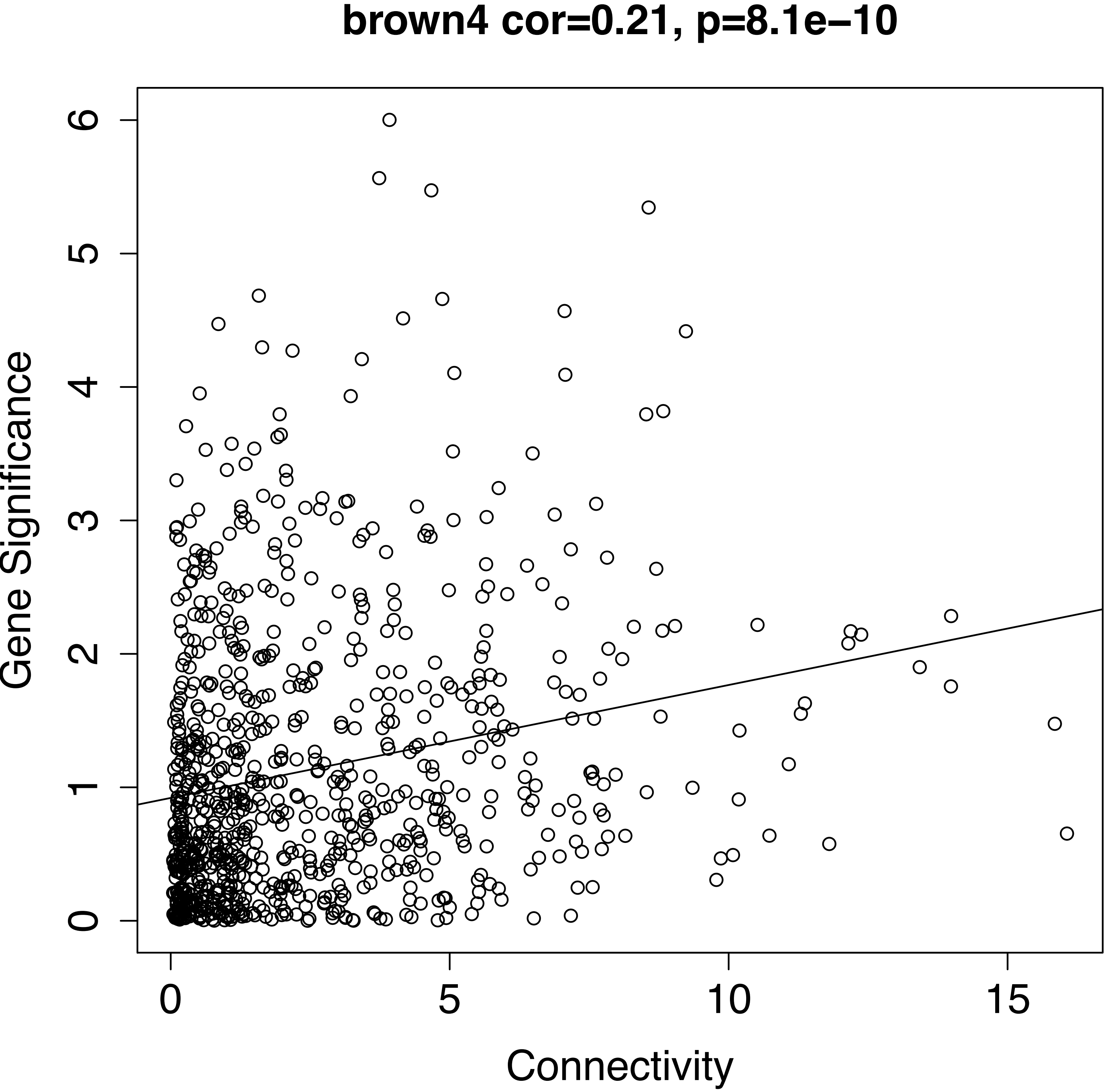

Supplementary
Figure 4 (a)

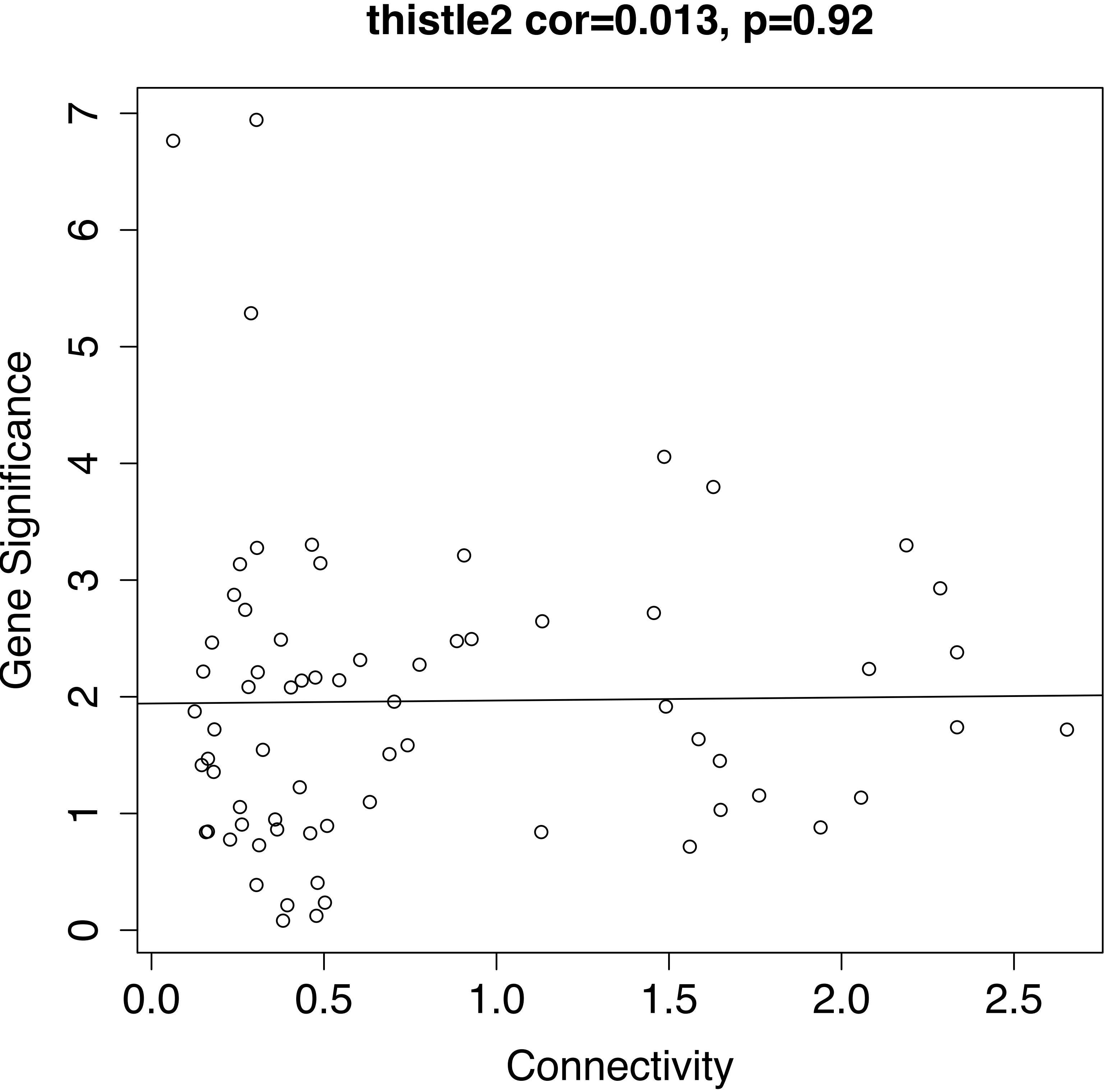

| Supp table 1: Differntialy expressed genes in brain of AD subjects at 25% FDR | | | | | | | |
|---|---|---|---|---|---|---|---|
| **ID** | **Gene** | **log2FoldCha** | **FC** | **lfcSE** | **pvalue** | **padj** | **Module Color** |
| ENSG00000138741 | TRPC3 | -0.29 | 0.82 | 0.05 | 4.57E-09 | 4.16E-05 | brown |
| ENSG00000196169 | KIF19 | 0.31 | 1.24 | 0.05 | 5.74E-09 | 4.16E-05 | blue |
| ENSG00000164946 | FREM1 | -0.26 | 0.83 | 0.05 | 1.14E-07 | 5.49E-04 | thistle2 |
| ENSG00000110723 | EXPH5 | -0.24 | 0.85 | 0.05 | 1.72E-07 | 6.22E-04 | thistle2 |
| ENSG00000166819 | PLIN1 | 0.29 | 1.22 | 0.06 | 6.58E-07 | 1.65E-03 | black |
| ENSG00000175455 | CCDC14 | -0.13 | 0.91 | 0.03 | 6.84E-07 | 1.65E-03 | paleturquoise |
| ENSG00000096060 | FKBP5 | 0.28 | 1.21 | 0.06 | 9.96E-07 | 1.80E-03 | brown4 |
| ENSG00000100433 | KCNK10 | 0.19 | 1.14 | 0.04 | 1.08E-06 | 1.80E-03 | blue |
| ENSG00000204950 | LRRC10B | 0.26 | 1.20 | 0.05 | 1.16E-06 | 1.80E-03 | grey60 |
| ENSG00000139144 | PIK3C2G | -0.28 | 0.82 | 0.06 | 1.32E-06 | 1.80E-03 | grey |
| ENSG00000162929 | KIAA1841 | -0.13 | 0.91 | 0.03 | 1.50E-06 | 1.80E-03 | brown |
| ENSG00000164604 | GPR85 | -0.22 | 0.86 | 0.05 | 1.54E-06 | 1.80E-03 | brown |
| ENSG00000007062 | PROM1 | -0.24 | 0.84 | 0.05 | 1.61E-06 | 1.80E-03 | brown |
| ENSG00000167772 | ANGPTL4 | 0.25 | 1.19 | 0.05 | 2.72E-06 | 2.81E-03 | brown4 |
| ENSG00000154721 | JAM2 | -0.19 | 0.88 | 0.04 | 2.91E-06 | 2.81E-03 | black |
| ENSG00000162654 | GBP4 | -0.26 | 0.83 | 0.06 | 3.94E-06 | 3.56E-03 | brown |
| ENSG00000111404 | RERGL | -0.26 | 0.83 | 0.06 | 4.50E-06 | 3.56E-03 | brown |
| ENSG00000142089 | IFITM3 | 0.27 | 1.20 | 0.06 | 4.52E-06 | 3.56E-03 | brown4 |
| ENSG00000106714 | CNTNAP3 | 0.18 | 1.14 | 0.04 | 4.90E-06 | 3.56E-03 | blue |
| ENSG00000183856 | IQGAP3 | 0.26 | 1.20 | 0.06 | 4.91E-06 | 3.56E-03 | grey |
| ENSG00000142449 | FBN3 | -0.26 | 0.84 | 0.06 | 5.47E-06 | 3.77E-03 | grey60 |
| ENSG00000153822 | KCNJ16 | -0.25 | 0.84 | 0.06 | 6.77E-06 | 4.45E-03 | black |
| ENSG00000170959 | DCDC1 | -0.23 | 0.85 | 0.05 | 1.01E-05 | 6.33E-03 | turquoise |
| ENSG00000117009 | KMO | -0.23 | 0.85 | 0.05 | 1.05E-05 | 6.33E-03 | mediumpurple3 |
| ENSG00000107562 | CXCL12 | -0.25 | 0.84 | 0.06 | 1.20E-05 | 6.95E-03 | brown |
| ENSG00000206432 | TMEM200C | -0.17 | 0.89 | 0.04 | 1.53E-05 | 8.54E-03 | lightsteelblue1 |
| ENSG00000138336 | TET1 | -0.14 | 0.91 | 0.03 | 1.70E-05 | 9.12E-03 | paleturquoise |
| ENSG00000172878 | METAP1D | -0.14 | 0.91 | 0.03 | 1.82E-05 | 9.43E-03 | turquoise |
| ENSG00000198830 | HMGN2 | 0.19 | 1.14 | 0.04 | 2.04E-05 | 9.98E-03 | paleturquoise |
| ENSG00000102359 | SRPX2 | 0.24 | 1.18 | 0.06 | 2.07E-05 | 9.98E-03 | brown4 |
| ENSG00000129353 | SLC44A2 | 0.18 | 1.14 | 0.04 | 2.19E-05 | 1.01E-02 | brown4 |
| ENSG00000198408 | MGEA5 | 0.12 | 1.09 | 0.03 | 2.23E-05 | 1.01E-02 | darkgrey |
| ENSG00000205221 | VIT | -0.23 | 0.85 | 0.06 | 2.50E-05 | 1.10E-02 | brown |
| ENSG00000141736 | ERBB2 | 0.23 | 1.18 | 0.06 | 2.70E-05 | 1.14E-02 | brown4 |
| ENSG00000183090 | FREM3 | -0.24 | 0.84 | 0.06 | 2.75E-05 | 1.14E-02 | mediumpurple3 |
| ENSG00000132470 | ITGB4 | 0.24 | 1.18 | 0.06 | 3.06E-05 | 1.21E-02 | brown4 |
| ENSG00000141314 | RHBDL3 | 0.18 | 1.13 | 0.04 | 3.10E-05 | 1.21E-02 | lightsteelblue1 |
| ENSG00000147872 | PLIN2 | 0.24 | 1.18 | 0.06 | 3.82E-05 | 1.46E-02 | brown4 |
| ENSG00000143341 | HMCN1 | -0.24 | 0.85 | 0.06 | 3.93E-05 | 1.46E-02 | lightgreen |
| ENSG00000111696 | NT5DC3 | 0.16 | 1.12 | 0.04 | 4.40E-05 | 1.55E-02 | lightsteelblue1 |
| ENSG00000124145 | SDC4 | 0.24 | 1.18 | 0.06 | 4.45E-05 | 1.55E-02 | black |
| ENSG00000214872 | SMTNL1 | 0.22 | 1.17 | 0.05 | 4.60E-05 | 1.55E-02 | turquoise |
| ENSG00000165899 | OTOGL | -0.20 | 0.87 | 0.05 | 4.61E-05 | 1.55E-02 | lightsteelblue1 |
| ENSG00000147647 | DPYS | -0.19 | 0.88 | 0.05 | 5.03E-05 | 1.59E-02 | lightsteelblue1 |
| ENSG00000074410 | CA12 | 0.21 | 1.16 | 0.05 | 5.04E-05 | 1.59E-02 | brown4 |
| ENSG00000116711 | PLA2G4A | -0.21 | 0.86 | 0.05 | 5.05E-05 | 1.59E-02 | brown |
| ENSG00000081248 | CACNA1S | -0.23 | 0.85 | 0.06 | 5.25E-05 | 1.61E-02 | grey60 |

| | | | | | | | |
|---|---|---|---|---|---|---|---|
| ENSG00000116717 | GADD45A | 0.24 | 1.18 | 0.06 | 5.34E-05 | 1.61E-02 | brown4 |
| ENSG00000120756 | PLS1 | -0.14 | 0.90 | 0.04 | 5.49E-05 | 1.62E-02 | lightsteelblue1 |
| ENSG00000048052 | HDAC9 | -0.15 | 0.90 | 0.04 | 5.62E-05 | 1.63E-02 | lightsteelblue1 |
| ENSG00000137857 | DUOX1 | 0.19 | 1.14 | 0.05 | 6.08E-05 | 1.71E-02 | yellow |
| ENSG00000113369 | ARRDC3 | 0.23 | 1.17 | 0.06 | 6.18E-05 | 1.71E-02 | brown4 |
| ENSG00000244694 | PTCHD4 | -0.16 | 0.89 | 0.04 | 6.26E-05 | 1.71E-02 | mediumpurple3 |
| ENSG00000050165 | DKK3 | 0.13 | 1.09 | 0.03 | 7.62E-05 | 2.04E-02 | brown |
| ENSG00000122863 | CHST3 | 0.23 | 1.17 | 0.06 | 7.85E-05 | 2.07E-02 | brown4 |
| ENSG00000187193 | MT1X | 0.22 | 1.16 | 0.06 | 8.09E-05 | 2.09E-02 | brown4 |
| ENSG00000154736 | ADAMTS5 | -0.19 | 0.88 | 0.05 | 8.23E-05 | 2.09E-02 | navajowhite2 |
| ENSG00000145681 | HAPLN1 | -0.21 | 0.87 | 0.05 | 8.78E-05 | 2.19E-02 | thistle2 |
| ENSG00000010322 | NISCH | 0.12 | 1.09 | 0.03 | 9.15E-05 | 2.21E-02 | yellow |
| ENSG00000188729 | OSTN | -0.23 | 0.85 | 0.06 | 9.15E-05 | 2.21E-02 | brown |
| ENSG00000158480 | SPATA2 | 0.12 | 1.09 | 0.03 | 9.68E-05 | 2.30E-02 | yellow |
| ENSG00000060656 | PTPRU | 0.16 | 1.11 | 0.04 | 0.00010754 | 2.51E-02 | yellow |
| ENSG00000110092 | CCND1 | -0.21 | 0.87 | 0.05 | 0.00011148 | 2.52E-02 | lightgreen |
| ENSG00000108551 | RASD1 | 0.21 | 1.15 | 0.05 | 0.00011168 | 2.52E-02 | brown4 |
| ENSG00000173011 | TADA2B | 0.10 | 1.07 | 0.03 | 0.00011371 | 2.52E-02 | yellow |
| ENSG00000124205 | EDN3 | -0.22 | 0.86 | 0.06 | 0.0001164 | 2.52E-02 | brown |
| ENSG00000164104 | HMGB2 | 0.21 | 1.16 | 0.06 | 0.00011701 | 2.52E-02 | brown4 |
| ENSG00000139304 | PTPRQ | -0.21 | 0.87 | 0.05 | 0.00011827 | 2.52E-02 | turquoise |
| ENSG00000131187 | F12 | 0.19 | 1.14 | 0.05 | 0.00012814 | 2.68E-02 | grey60 |
| ENSG00000128578 | STRIP2 | -0.13 | 0.91 | 0.03 | 0.00012947 | 2.68E-02 | mediumpurple3 |
| ENSG00000141469 | SLC14A1 | 0.21 | 1.16 | 0.06 | 0.00013691 | 2.79E-02 | black |
| ENSG00000155287 | SLC25A28 | 0.10 | 1.07 | 0.03 | 0.00014467 | 2.90E-02 | blue |
| ENSG00000144908 | ALDH1L1 | 0.22 | 1.16 | 0.06 | 0.00015171 | 2.90E-02 | brown4 |
| ENSG00000183323 | CCDC125 | -0.11 | 0.93 | 0.03 | 0.00015593 | 2.90E-02 | blue |
| ENSG00000162367 | TAL1 | -0.21 | 0.86 | 0.06 | 0.00015631 | 2.90E-02 | lightcyan |
| ENSG00000175104 | TRAF6 | 0.10 | 1.07 | 0.03 | 0.00015828 | 2.90E-02 | darkgrey |
| ENSG00000122574 | WIPF3 | 0.15 | 1.11 | 0.04 | 0.00015889 | 2.90E-02 | yellow |
| ENSG00000171189 | GRIK1 | -0.16 | 0.89 | 0.04 | 0.00015929 | 2.90E-02 | thistle2 |
| ENSG00000102362 | SYTL4 | 0.22 | 1.16 | 0.06 | 0.00016008 | 2.90E-02 | brown4 |
| ENSG00000151929 | BAG3 | 0.21 | 1.16 | 0.06 | 0.00016021 | 2.90E-02 | brown4 |
| ENSG00000152818 | UTRN | -0.14 | 0.91 | 0.04 | 0.00016545 | 2.93E-02 | lightgreen |
| ENSG00000111907 | TPD52L1 | 0.21 | 1.16 | 0.06 | 0.00016585 | 2.93E-02 | black |
| ENSG00000138246 | DNAJC13 | -0.09 | 0.94 | 0.02 | 0.0001791 | 3.12E-02 | darkgrey |
| ENSG00000123473 | STIL | -0.12 | 0.92 | 0.03 | 0.00019321 | 3.33E-02 | brown |
| ENSG00000101413 | RPRD1B | 0.08 | 1.05 | 0.02 | 0.00019661 | 3.34E-02 | brown4 |
| ENSG00000141682 | PMAIP1 | -0.22 | 0.86 | 0.06 | 0.00019847 | 3.34E-02 | navajowhite2 |
| ENSG00000118777 | ABCG2 | -0.21 | 0.87 | 0.06 | 0.00021336 | 3.55E-02 | brown |
| ENSG00000120907 | ADRA1A | -0.16 | 0.90 | 0.04 | 0.00022169 | 3.63E-02 | salmon4 |
| ENSG00000178015 | GPR150 | 0.18 | 1.13 | 0.05 | 0.00022565 | 3.63E-02 | brown |
| ENSG00000058866 | DGKG | 0.15 | 1.11 | 0.04 | 0.00022725 | 3.63E-02 | brown4 |
| ENSG00000174576 | NPAS4 | 0.12 | 1.09 | 0.03 | 0.00023018 | 3.63E-02 | darkslateblue |
| ENSG00000225828 | FAM229A | 0.19 | 1.14 | 0.05 | 0.00023524 | 3.63E-02 | yellow |
| ENSG00000112706 | IMPG1 | -0.16 | 0.89 | 0.04 | 0.00023571 | 3.63E-02 | darkgrey |
| ENSG00000103742 | IGDCC4 | 0.19 | 1.14 | 0.05 | 0.00023789 | 3.63E-02 | brown4 |
| ENSG00000136297 | MMD2 | -0.21 | 0.86 | 0.06 | 0.00024055 | 3.63E-02 | salmon4 |
| ENSG00000166035 | LIPC | -0.20 | 0.87 | 0.06 | 0.00024097 | 3.63E-02 | grey |
| ENSG00000173085 | COQ2 | -0.15 | 0.90 | 0.04 | 0.00024339 | 3.63E-02 | paleturquoise |

| | | | | | | | |
|---|---|---|---|---|---|---|---|
| ENSG00000175344 | CHRNA7 | -0.13 | 0.91 | 0.05 | 0.01334687 | 2.42E-01 | thistle2 |
| ENSG00000138185 | ENTPD1 | -0.07 | 0.95 | 0.03 | 0.01337397 | 2.42E-01 | paleturquoise |
| ENSG00000187079 | TEAD1 | 0.10 | 1.07 | 0.04 | 0.01337623 | 2.42E-01 | black |
| ENSG00000138074 | SLC5A6 | -0.12 | 0.92 | 0.05 | 0.01339205 | 2.42E-01 | darkolivegreen |
| ENSG00000188486 | H2AFX | 0.10 | 1.07 | 0.04 | 0.01343747 | 2.43E-01 | yellow |
| ENSG00000182551 | ADI1 | 0.10 | 1.07 | 0.04 | 0.01349617 | 2.43E-01 | black |
| ENSG00000133878 | DUSP26 | 0.08 | 1.06 | 0.03 | 0.01350653 | 2.43E-01 | darkolivegreen |
| ENSG00000186591 | UBE2H | 0.06 | 1.04 | 0.02 | 0.0135286 | 2.43E-01 | brown4 |
| ENSG00000075240 | GRAMD4 | -0.08 | 0.95 | 0.03 | 0.01354124 | 2.43E-01 | grey60 |
| ENSG00000102878 | HSF4 | 0.11 | 1.08 | 0.05 | 0.01357758 | 2.44E-01 | tan |
| ENSG00000197372 | ZNF675 | -0.08 | 0.95 | 0.03 | 0.01365746 | 2.44E-01 | brown |
| ENSG00000086544 | ITPKC | 0.13 | 1.10 | 0.05 | 0.01367243 | 2.44E-01 | brown4 |
| ENSG00000185973 | TMLHE | -0.08 | 0.94 | 0.03 | 0.01368991 | 2.44E-01 | paleturquoise |
| ENSG00000149480 | MTA2 | 0.11 | 1.08 | 0.05 | 0.01370395 | 2.44E-01 | brown4 |
| ENSG00000155085 | AK9 | -0.08 | 0.95 | 0.03 | 0.01371475 | 2.44E-01 | paleturquoise |
| ENSG00000145700 | ANKRD31 | -0.12 | 0.92 | 0.05 | 0.01371854 | 2.44E-01 | blue |
| ENSG00000058335 | RASGRF1 | -0.11 | 0.93 | 0.04 | 0.01375805 | 2.45E-01 | grey60 |
| ENSG00000166596 | WDR16 | -0.11 | 0.93 | 0.04 | 0.01387111 | 2.46E-01 | lightsteelblue1 |
| ENSG00000102678 | FGF9 | -0.09 | 0.94 | 0.04 | 0.01387644 | 2.46E-01 | saddlebrown |
| ENSG00000181938 | GINS3 | -0.11 | 0.92 | 0.05 | 0.01391095 | 2.46E-01 | blue |
| ENSG00000168724 | DNAJC21 | -0.07 | 0.95 | 0.03 | 0.01392879 | 2.46E-01 | brown |
| ENSG00000165912 | PACSIN3 | -0.13 | 0.91 | 0.05 | 0.01399602 | 2.47E-01 | blue |
| ENSG00000138792 | ENPEP | 0.14 | 1.10 | 0.06 | 0.01400196 | 2.47E-01 | darkslateblue |
| ENSG00000188739 | RBM34 | 0.10 | 1.07 | 0.04 | 0.01401802 | 2.47E-01 | yellow |
| ENSG00000179476 | C14orf28 | -0.09 | 0.94 | 0.04 | 0.01403361 | 2.47E-01 | paleturquoise |
| ENSG00000137275 | RIPK1 | 0.09 | 1.06 | 0.04 | 0.01404179 | 2.47E-01 | brown4 |
| ENSG00000188677 | PARVB | 0.08 | 1.06 | 0.03 | 0.0140672 | 2.47E-01 | darkolivegreen |
| ENSG00000258839 | MC1R | 0.13 | 1.10 | 0.05 | 0.01410173 | 2.47E-01 | yellow |
| ENSG00000143669 | LYST | -0.06 | 0.96 | 0.02 | 0.01417656 | 2.48E-01 | lightsteelblue1 |

ID: Ensembl id of the gene
log2Fold change: log2 value of the fold change calculated using DeSeq2
FC: Fold change
pvalue: P value for the fold change of expression among AD and controls
padj: FDR adjusted pvalue
ModuleColor: The group assigned to each gene on the basis of co-expression by WGCNA

**Supp table 2a: Differntialy expressed hub genes of Thistle2 module in brain of AD subjects**

| id | gene | log2FoldChange | lfcSE | pvalue | Mouse/ Rat brain (Diff expression P) | Region | Ethanol/ Treatment | Refernce |
|---|---|---|---|---|---|---|---|---|
| ENSG00000171189 | GRIK1 | -0.16 | 0.04 | 1.59E-04 | 3.00E-02 | Frontal-cortex | AP vs nAP | Ferguson et al, 2017 |
| ENSG00000189056 | RELN | -0.20 | 0.06 | 5.04E-04 | 6.80E-03 | whole brain | AP vs nAP | Kimpel et al, 2007 |
| ENSG00000231764 | DLX6-AS1 | -0.18 | 0.06 | 1.18E-03 | | | | |
| ENSG00000146469 | VIP | -0.16 | 0.06 | 4.16E-03 | 9.00E-04 | Whole brain | AP vs nAP | Mulligan et al, 2006 |
| ENSG00000157404 | KIT | -0.14 | 0.05 | 5.77E-03 | 4.00E-02 | Central Amygdala | AP vs nAP | Ferguson et al, 2017 |
| ENSG00000017427 | IGF1 | -0.12 | 0.05 | 1.91E-02 | | | | |
| ENSG00000135824 | RGS8 | -0.13 | 0.06 | 2.31E-02 | | | | |
| ENSG00000172137 | CALB2 | -0.12 | 0.06 | 3.54E-02 | 1.90E-02 | Frontal cortex | DID Drinking | Osterndorff-Kahanek et al, 2013 |
| ENSG00000178568 | ERBB4 | -0.06 | 0.04 | 9.32E-02 | | | | |
| ENSG00000144355 | DLX1 | -0.08 | 0.05 | 1.32E-01 | 4.90E-03 | whole brain | AP vs nAP | Mulligan et al, 2006 |
| ENSG00000151136 | BTBD11 | -0.06 | 0.05 | 1.93E-01 | 2.00E-02 | Frontal-cortex | Every other day drinking | Osterndorff-Kahanek et al, 2013 |

ID: Ensembl id of the gene
log2Fold change: log2 value of the fold change calculated using DeSeq2
Column F: Diffential expression P value in mouse or rat models
Column G: Region of the mouse/ rat brain used in the study
Column H: Description of the mouse model (e.g. Alcohol prefernce vs non prefernce; DID (Drinking in the dark)
Reference: Reference of the study

**Supp table 2b: Differntialy expressed hub genes of Brown 4 module in brain of AD subjects**

| id | gene | log2FoldChar | lfcSE | pvalue | **Mouse/ Rat brain (Diff expression P)** | **Region** | **Ethanol/ Treatment** | **Refernce** |
|---|---|---|---|---|---|---|---|---|
| ENSG00000142089 | IFITM3 | 0.27 | 0.06 | 4.52E-06 | 7.70E-04 | Frontal-cortex | AP vs nAP | Osterndorff-Kahanek et al, 2013 |
| ENSG00000147872 | PLIN2 | 0.24 | 0.06 | 3.82E-05 | | | | |
| ENSG00000144908 | ALDH1L1 | 0.22 | 0.06 | 1.52E-04 | 1.90E-02 | Frontal-cortex | Chronic drinking | Osterndorff-Kahanek et al, 2013 |
| ENSG00000151929 | BAG3 | 0.21 | 0.06 | 1.60E-04 | | | | |
| ENSG00000148498 | PARD3 | 0.17 | 0.05 | 1.21E-03 | | | | |
| ENSG00000185201 | IFITM2 | 0.17 | 0.06 | 1.90E-03 | 1.20E-03 | Frontal-cortex | AP vs nAP | Osterndorff-Kahanek et al, 2013 |
| ENSG00000072952 | MRVI1 | 0.17 | 0.06 | 2.30E-03 | | | | |
| ENSG00000067182 | TNFRSF1A | 0.16 | 0.06 | 5.20E-03 | | | | |
| ENSG00000077238 | IL4R | 0.16 | 0.06 | 6.06E-03 | | | | |
| ENSG00000051620 | HEBP2 | 0.13 | 0.05 | 6.18E-03 | | | | |
| ENSG00000109113 | RAB34 | 0.15 | 0.06 | 6.26E-03 | 7.20E-03 | Frontal-cortex | AP vs nAP | Osterndorff-Kahanek et al, 2013 |
| ENSG00000168309 | FAM107A | 0.15 | 0.05 | 6.71E-03 | 1.60E-03 | Whole Brain | AP vs nAP | Mulligan et al, 2006 |
| ENSG00000145623 | OSMR | 0.15 | 0.06 | 6.76E-03 | 7.70E-03 | Whole Brain | AP vs nAP | Mulligan et al, 2006 |
| ENSG00000197324 | LRP10 | 0.14 | 0.05 | 7.17E-03 | | | | |
| ENSG00000155366 | RHOC | 0.14 | 0.05 | 8.35E-03 | 4.50E-05 | Whole Brain | AP vs nAP | Mulligan et al, 2006 |
| ENSG00000187091 | PLCD1 | 0.13 | 0.05 | 9.15E-03 | | | | |
| ENSG00000183255 | PTTG1IP | 0.13 | 0.05 | 1.26E-02 | 4.30E-04 | Frontal-cortex | AP vs nAP | Osterndorff-Kahanek et al, 2013 |
| ENSG00000117519 | CNN3 | 0.13 | 0.06 | 1.75E-02 | 1.20E-02 | Whole Brain | AP vs nAP | Kimpel et al, 2007 |
| ENSG00000182541 | LIMK2 | 0.12 | 0.05 | 2.03E-02 | | | | |
| ENSG00000158710 | TAGLN2 | 0.13 | 0.06 | 2.35E-02 | 2.80E-02 | Frontal-cortex | AP vs nAP | Ferguson et al, 2017 |
| ENSG00000148175 | STOM | 0.12 | 0.06 | 2.81E-02 | | | | |
| ENSG00000168610 | STAT3 | 0.10 | 0.05 | 3.34E-02 | 7.00E-04 | Whole Brain | AP vs nAP | Mulligan et al, 2006 |
| ENSG00000135926 | TMBIM1 | 0.12 | 0.06 | 3.75E-02 | 3.00E-03 | Whole Brain | AP vs nAP | Mulligan et al, 2006 |
| ENSG00000131981 | LGALS3 | 0.11 | 0.06 | 6.71E-02 | | | | |
| ENSG00000129116 | PALLD | 0.08 | 0.05 | 9.48E-02 | | | | |
| ENSG00000198805 | PNP | 0.10 | 0.06 | 1.01E-01 | | | | |
| ENSG00000181826 | RELL1 | 0.09 | 0.06 | 1.23E-01 | 4.20E-02 | Frontal-cortex | DID drinking | Osterndorff-Kahanek et al, 2013 |
| ENSG00000119408 | NEK6 | 0.06 | 0.05 | 1.63E-01 | 1.20E-02 | Whole Brain | AP vs nAP | Mulligan et al, 2006 |
| ENSG00000137693 | YAP1 | 0.07 | 0.05 | 1.73E-01 | | | | |
| ENSG00000031081 | ARHGAP31 | 0.06 | 0.05 | 2.09E-01 | | | | |
| ENSG00000147065 | MSN | 0.07 | 0.05 | 2.22E-01 | 5.00E-03 | Whole Brain | AP vs nAP | Mulligan et al, 2006 |
| ENSG00000077150 | NFKB2 | 0.07 | 0.06 | 2.27E-01 | | | | |
| ENSG00000162909 | CAPN2 | 0.04 | 0.04 | 2.31E-01 | 1.10E-02 | Frontal-cortex | AP vs nAP | Osterndorff-Kahanek et al, 2013 |
| ENSG00000177119 | ANO6 | 0.06 | 0.05 | 2.34E-01 | | | | |
| ENSG00000106565 | TMEM176B | 0.07 | 0.06 | 2.55E-01 | 2.30E-02 | Frontal-cortex | AP vs nAP | Osterndorff-Kahanek et al, 2013 |
| ENSG00000213719 | CLIC1 | 0.06 | 0.06 | 2.66E-01 | | | | |
| ENSG00000168994 | PXDC1 | 0.06 | 0.06 | 3.04E-01 | | | | |
| ENSG00000101608 | MYL12A | 0.04 | 0.05 | 4.93E-01 | | | | |
| ENSG00000124782 | RREB1 | 0.02 | 0.05 | 6.95E-01 | 4.70E-02 | Frontal-cortex | AP vs nAP | Osterndorff-Kahanek et al, 2013 |

ID: Ensembl id of the gene
log2Fold change: log2 value of the fold change calculated using DeSeq2
Column F: Diffential expression P value in mouse or rat models
Column G: Region of the mouse/ rat brain used in the study
Column H: Description of the mouse model (e.g. Alcohol prefernce vs non prefernce; DID (Drinking in the dark)
Reference: Reference of the study
